# Phenotypic plasticity of ER+ breast cancer in the bone microenvironment

**DOI:** 10.1101/2020.11.14.383000

**Authors:** Igor L. Bado, Hai Wang, Poonam Sarkar, Jun Liu, William Wu, Weijie Zhang, Hin-Ching Lo, Aaron Muscallera, Mahnaz Janghorban, Ik-Sun Kim, Swarnima Singh, Amit Goldstein, Purba Singh, Huang Shixia, Gaber M. Waleed, Matthew J. Ellis, Xiang H.-F. Zhang

## Abstract

ER+ breast cancer exhibits a strong bone-tropism in metastasis. How the bone microenvironment impacts the ER signaling and endocrine therapies remains poorly understood. Here, we discover that the osteogenic niche transiently reduces ER expression and activities specifically in bone micrometastases (BMMs), leading to endocrine resistance. This is mediated by gap junctions and paracrine FGF/PDGF signaling, which together generate a stable “memory”: cancer cells extracted from bone remain resistant to endocrine therapies for several generations. Using single cell-derived populations (SCPs), we demonstrated that this process is independent of clonal selection, and represents an EZH2-mediated epigenomic reprogramming. EZH2 drives ER+ BMMs toward a basal and stem-like state. EZH2 inhibition reverses endocrine resistance. Our data demonstrates how epigenomic adaptation to the bone microenvironment drives phenotypic plasticity of metastatic seeds and alters their therapeutic responses together with clonal selection, and provides insights into the clinical enigma of ER+ metastatic recurrences despite endocrine therapies.

## INTRODUCTION

Estrogen receptor positive (ER+) breast cancer accounts for over 70% of all breast cancers, and after recurring, causes over 24,000 deaths per year in the US. Adjuvant endocrine therapies target ER and significantly reduce metastatic recurrences. However, 20-40% of patients still develop metastases, often after a prolonged latency (Lim et al., 2012; Zhang et al., 2013). Thus, it is imperative to understand how disseminated ER+ cancer cells escape endocrine therapies in distant organs and to identify therapies that can eliminate these cells.

Bone is the most frequently affected organ by ER+ breast cancer, which is usually luminal-like. Compared to the basal-like subtype, luminal A/B breast cancer exhibits a 2.5-fold increased frequency of bone metastasis, but a 2.5-fold decreased frequency of lung metastasis (Kennecke et al., 2010; Smid et al., 2008). Bone metastases of luminal-like breast cancer are usually late-onset, occurring beyond 5 years after surgery. Current tumor-intrinsic biomarkers based on primary tumors can predict recurrences within 5 years, but cannot predict late-onset recurrences after 5 years (Sgroi et al., 2013), suggesting that the capacity of developing late-onset metastasis is not encoded in cancer cells. Thus, there appear to be unique interactions between the bone microenvironment and ER+ disseminated tumor cells that allow them to survive adjuvant endocrine therapies and persist for a prolonged time.

Very little is known about how the bone microenvironment affects ER+ breast cancer cells in terms of key signaling pathways (e.g., ER signaling itself), therapeutic responses, and the evolution process. Seemingly, conflicting findings were reported, suggesting profound uncharacterized biology. For instance, a few studies revealed a paradoxically high disconcordance rate of ER status between primary tumors and DTCs, suggesting loss of ER in DTCs, which may be related to resistance to endocrine therapies (Fehm et al., 2008; Jäger et al., 2015). On the other hand, it was also noted that most clinically-detectable macroscopic bone metastases (>85%) remain positive for ER (Hoefnagel et al., 2013), seemingly contradicting the DTC findings. ER+ bone metastases still respond to endocrine therapies in many cases, although resistance almost invariably develops. These clinical observations cry for mechanistic studies, which are hindered by the lack of ER+ bone metastasis models, as well as technical limitations of detecting/investigating metastatic cells at a single cell resolution.

We have previously developed a series of models and techniques to investigate cancer-bone interaction at a single cell resolution. We discovered that bone micrometastases are usually localized in close contact with osteogenic cells including mesenchymal stem cells (MSCs) and osteoblasts (OBs)(Wang et al., 2015). Direct interaction between cancer and osteogenic cells is mediated by adherens and gap junctions, which stimulate mTOR and calcium signaling in cancer cells, respectively (Wang et al., 2015, 2018). This interaction precedes the onset of osteolytic vicious cycle that has been well characterized before, and represents an intermediate stage of bone colonization between dormant single DTCs and overt macrometastases. Herein, by combining our techniques with an array of ER+ breast cancer models, we aimed to understand how the bone microenvironment may dictate the evolution of ER+ breast cancer cells, leading to unexpected cellular alterations and therapeutic responses.

## RESULTS

### Microscopic bone lesions transiently lose ER expression

To study how ER+ breast cancer cells interact with the bone microenvironment, we identified two patient-derived xenograft models (PDXs), HCI011 and WHIM9, which were developed from patients with bone metastases (DeRose et al., 2011; Li et al., 2013) and exhibited spontaneous metastasis from mammary glands to bone in immunodeficient mice 5-6 months after orthotopic tumors are removed (Figure 1A). When we attempted to use human-specific ER as a marker to distinguish ER+ metastases from bone cells, we noticed that, spontaneous bone metastases exhibited reduced ER expression in both PDX models (Figure 1B). In WHIM9, the sizes of bone metastases are diverse and display an interesting correlation with intensity of nuclear ER by IHC staining (Figure S1B). Moreover, in macroscopic bone metastases, ER expression appeared to be heterogeneous: cancer cells adjacent to bone matrix seemed to exhibit weaker IHC staining of ER (Figure S1C). These observations raised interesting questions about the influence of the bone microenvironment on ER expression in ER+ breast cancer cells.

**Figure 1:**
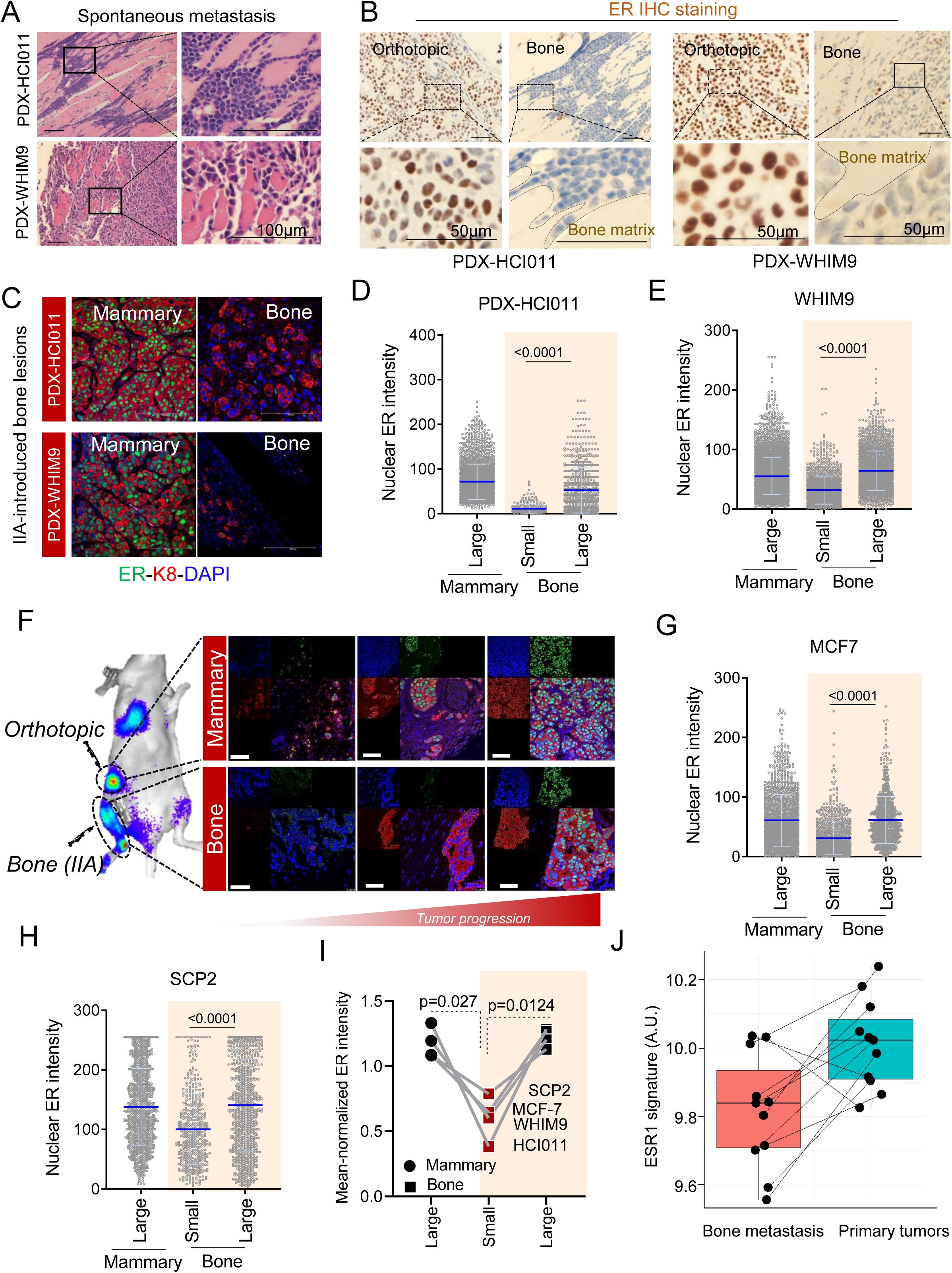
The bone microenvironment induces transient loss of ER expression in ER+ breast cancer cells. **A.** Representative H&E staining of spontaneous metastases of HCI011 and WHM9 tumors to spine and hind limb, respectively. Scale bar: 100μm. **B.** Human-specific ER IHC staining are shown for spontaneous metastasis of HCI011 and WHIM9, respectively. Bone matrix is annotated. Scale bar: 50μm. **C.** Confocal images showing immunofluorescence (IF) staining of ER (green), keratin 8 [k8] (red), and DAPI (blue) in orthotopic (mammary) and IIA-induced bone metastasis models of ER+ PDXs (HCI011 and WHIM9). Scale bars: 100µM. Representative images were captured with a 40x oil objective lens **D.** Dot plot depicting quantification of nuclear ER intensity from HCI011 primary tumor and bone metastasis specimens as illustrated in Figure 1A. Bone lesions were classified into “small” and “big” groups based on cell numbers captured by a same field with the cutoffs being < median – 0.5xS.D. (small) or > median + 0.5xS.D. (big). Each dot represents the fluorescence intensity of ER of a single cells. Cells from 3-5 different animals are plotted. **E.** Dot plot depicting quantification of nuclear ER intensity from WHIM9 primary tumor and bone metastasis specimens as illustrated in Figure 1A. Bone lesions were classified into “small” and “big” groups as defined in (B). Each dot represents the fluorescence intensity of ER of a single cells. Cells from 3-5 different animals are plotted. **F.** Representative IF images of MCF7 cells following orthotopic and bone transplantation in nude mice. Changes in ER expression are illustrated in primary tumor and bone metastasis at different stages of tumor progression. Early, intermediate and late phases are depicted from left to right. Green, red and blue represent IF staining of estrogen receptor (ER), cytokeratin (K8) and nucleus (DAPI). Scale bars, 50μm. **G.** Dot plot depicting quantification of nuclear ER intensity from MCF7 primary tumor and bone metastasis specimens as illustrated in (D). Bone lesions were classified into “small” and “big” as defined in (B). Each dot represents the fluorescence intensity of ER of a single cells. Cells from 6 different animals are plotted. **H.** Dot plot depicting quantification of nuclear ER intensity from MCF7 single cell-derived population 2 (SCP2) primary tumor and bone metastasis specimens. Bone lesions were classified into “small” and “big” groups as defined in (B). Each dot represents the fluorescence intensity of ER of a single cells. Cells from 4 different animals are plotted. **I.** Dot plot showing the mean-normalized ER intensity of all cancer models using from Figure 1A to 1F. P-values derive from a two-tailed paired Student’s *t*-test. **J.** Boxplot showing changes in ESR1 early signature in matched bone metastases and primary specimens from breast cancer patients ((https://github.com/npriedig/). All images were captured with Leica TCS SP5 confocal microscope. A 40x or 63x oil objective lens were used to capture all images (Immersion oil refractive index n=1.51). All quantifications were performed using ImageJ (Fiji). All statistical analyses represent a two-tailed unpaired Student’s *t*-test except when specified otherwise.

To further pursue the impact of bone microenvironment on ER+ breast cancer cells, we performed intra-iliac artery (IIA) injection of dissociated PDX cells to introduce experimental bone metastases to investigate early-stage bone colonization (Figure S1C). This approach synchronizes the onset of colonization and enriches microscopic metastases, thereby allowing quantitative examination of bone colonization of relatively indolent cancer cells at different temporal stages (Yu et al., 2016).

Like in spontaneous bone metastases, a strong correlation was found between ER expression and IIA-introduced lesion size in bone (Figure S1D) but not in orthotopic tumors (Figure S1E). When we classified tumors as micrometastasis (small) and macrometastasis (large) by cell counts, the expression of ER appeared to diminish in microscopic bone lesions compared to mammary tumors of the same models (Figure 1C) but was restored as bone lesions further progress into macroscopic bone metastases. (Figure 1D-E). We next asked if this phenomenon could also be observed in other models that are more amenable to experimental manipulations. MCF-7 cells in orthotopic tumors and IIA-introduced bone lesions from the same host animals were compared at different time points. Similar to PDXs, a loss of ER expression was found at early-stage of bone colonization (Figure 1F-G).

Two possible mechanisms might explain the differences in ER expression between bone lesions of different sizes. First, genetically distinct ER-low and ER-high cancer cell clones may pre-exist, and the former progresses at a much slower rate and only form small lesions. Second, there may be a transient and reversible loss of ER in ER+ cancer cells in the bone microenvironment. To distinguish these possibilities, we collected single cell-derived populations (SCPs) from MCF-7 cells. A SCP was expanded from a single cell, and therefore, is much more homogeneous genetically. Four SCPs were developed with variable tumorigenicity in mammary glands or bones (Figure S1F), suggesting that there is indeed pre-existing clonal heterogeneity as previously shown in other cell lines (Minn et al., 2005a). Using an SCP that exhibits strong tumorigenicity in both mammary glands and bone, we compared the ER expression between micro- and macro-metastases and observed the same difference as for parental MCF7 cells (Figure 1H). Taken together, data obtained from multiple models supported that the bone microenvironment induces a loss of ER expression specifically in micrometastases (Figure 1I).

The loss of ER was also observed on clinical specimens in previous studies (Hirata et al., 2009; Schrijver et al., 2018). In particular, in a study comparing gene expression profiles between patient-matched primary and bone metastases, ESR1 was found to be one of the top genes down-regulated in bone metastases (Priedigkeit et al., 2017). Gene Set Variation Analysis (GSVA) further suggests that there is an even stronger down-regulation of acute ER signaling (Figure 1J). Together with the experimental observations, this data provide an explanation for the discrepant findings in previous literatures, arguing that early-stage colonization of ER+ breast cancer cells in the bone is associated with a transient and reversible down-regulation of ER.

### Direct interaction with osteogenic cells mediates the loss of ER expression

Early-stage bone colonization of ER+ breast cancer cells frequently occurs in the niche comprised of cells with osteogenic capacity, including MSCs and OBs (Wang et al., 2015). Among MCF7-derived bone lesions in the same animal, there appear to be an inverse correlation between nuclear intensity of ER (assessed by immunofluorescence) and the abundance of osteogenic cells in the microenvironment niche (assessed by frequency of stromal cells expressing alkaline phosphatase [ALP]) (Figure 2A). Similar observations were found in IIA-induced bone metastasis models of Patient-derived xenografts, HCI011 and WHIM9 (Figure 2B-C), suggesting the interaction of cancer cells with osteogenic cells may provoke the transient loss of ER. To further dissect this interaction, we employed a fetal osteoblast cell line (FOB) and a human mesenchymal cell line (MSC) to represent osteogenic cells. Luminal-like cancer cells and osteogenic cells form heterotypic organoids in 3D suspension co-culture, which successfully recapitulated several aspects of cancer-niche interaction (Wang et al., 2015). Co-staining of ER and keratin 8 (K8, a marker of luminal cancer cells) in 3D co-cultures revealed a loss of ER in MCF-7 and HCI011 primary cells (Figure 2D) similar to *in vivo* bone micrometastases, suggesting that interaction with osteogenic cells in vitro can recapitulate the ER down-regulation in bone micrometastasis (BMMs).

**Figure 2:**
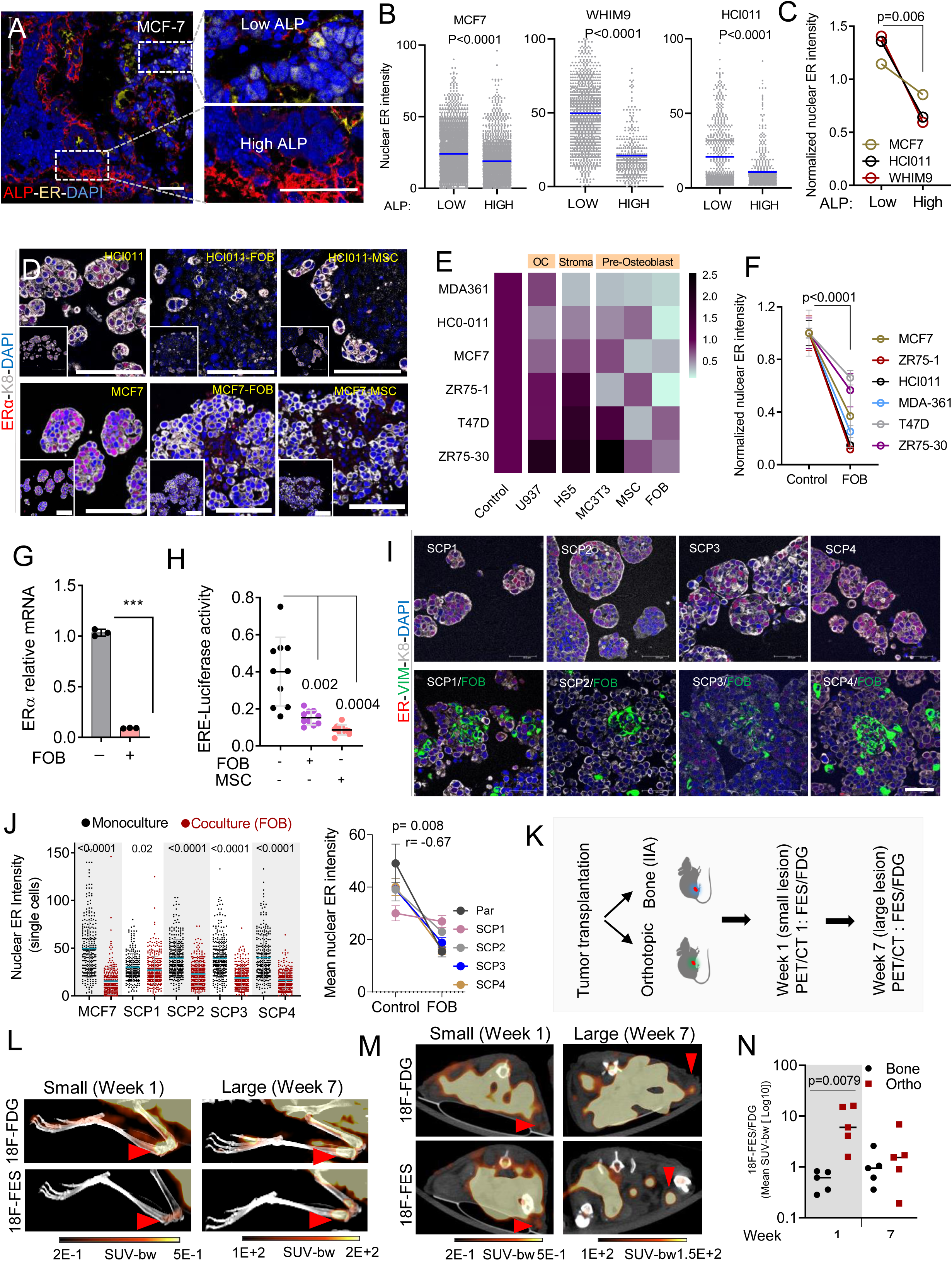
Osteogenic cells promote loss of ER expression and reduction of ER activities during early stages of bone colonization **A.** Representative confocal image of bone metastasis showing a negative association between ER expression in cancer cells and alkaline phosphatase (ALP) expression in osteogenic cells. ER, ALP and nuclei (DAPI) are shown as yellow, red, and blue, respectively. Scale bars : 50µM **B.** Dot plot showing quantification of ER expression in MCF7 and two PDX (HCI011 and WHIM9) models of IIA-induced bone metastasis. Alkaline phosphatase (ALP) was used to identify osteogenic cells (red). Bone metastases were classified as Low or High according to ALP expression in surrounding osteogenic cells as depicted in Figure 2A. Each dot represents a cell, and 3-5 animals were combined. **C.** Graph representing the paired analysis of ER expression according to alkaline phosphatase (ALP) enrichment in the tumor microenvironment as shown in A. Mean expression of ER was obtained by averaging single cell data from 3-5 different mice. P-values derive from two-tailed paired Student’s *t*-test. **D.** Representative IF images of HCI011-derived primary cells and MCF7 cells in 3D monoculture and co-culture with human fetal osteoblast cell line (FOB) and mesenchymal stem cell line (MSC). ER, keratin 8 (K8) and nuclei (DAPI) are represented in red, grey, and blue, respectively. Scale bars: 100µm. **E.** Heatmap showing the mean intensity of ER in primary cells (HCI011) and breast cancer cell lines (MDA-MB-361, MCF7, ZR75-1, T47D, ZR75-30) in 3D monoculture (control) or co-culture with osteoclast precursors (U937), bone marrow stromal cells (Hs5), mouse pre-osteoblasts (MC3T3), human mesenchymal stem cells (MSC) and human pre-osteoblast (FOB). All co-cultures were performed in triplicate and images were captured with a 40x oil objective lens. **F.** Graph comparing ER expression in monoculture versus co-culture of multiple cell lines with FOB. P-value results from a two-tailed paired Student’s *t*-test. 3 separated experiments were used. Error bars: mean +/- standard deviation. **G.** Relative mRNA expression of ESR1 in 3D monoculture or co-culture of MCF7 with FOB. Data result from MCF7 cells only (FACS sorted). **H.** Dot plot representing ER transcriptional activity in MCF7 cells expressing pGL2 ERE-luciferase reporter. MCF7 cells were cultured in 3D with or without osteogenic cells (FOB and MSC) for 7 days. Luciferase activity was assessed using IVIS Lumina II *in vivo* system; n=10 technical replicates. Error bars: mean +/- standard deviation. **I.** Representative confocal images showing ER expression (red) in MCF7 single cell-derived populations (SCPs) in 3D monoculture or co-culture with FOB cells. Vimentin (VIM), Keratin 8 (CK8), and DAPI were used to identify osteoblasts (Green), cancer cells (grey) and cell nuclei (blue), respectively. Scale bars: 50µm. **J.** Quantified ER expression from confocal images of single cell-derived populations (SCP1-SCP4). A two-tailed paired Student’s *t*-test analysis reveals a strong negative correlation (r) between monoculture and FOB co-cultures (right panel n=5). Error bars: +/- standard error of the mean. Each dot represents a cell from 3 different images. **K.** Diagram showing the experimental design for positron emission tomography–computed tomography (PET-CT) imaging of MCF7 cells transplanted orthotopically or to bone via IIA injection. Two rounds of imaging were performed at week 1 and week 5 post transplantation using 18F-Fluoroestradiol (18F-FES) and 18F-Fluorodeoxyglucose (18F-FDG) with 2 days apart. **L.** Representative PET/CT scans showing the maximum intensity projection (MIP) visualization of radiolabeled 18F-Fluorodeoxyglucose (18F-FDG) and 18F-Fluoroestradiol (18F-FES) in bone. Early time point (Week 1) and late time point (Week 7) were used to depict the micro-metastasis stage (small) and the macrometastasis stage (large). MCF7 bone metastases were generated using Intra-iliac artery injection. Red arrows indicate tumor location (Joint area). A smaller scale (0.2-0.5 SUV-bw) was used for week 1 images to allow detection of small lesions while a scale of 100-200 SUV-bw was used for the macrometastasis stage. **M.** Axial view of representative PET/CT scans depicting the uptake of radiolabeled fluorodeoxyglucose (18F-FDG) and fluoroestradiol (18F-FES) in small and large lesions of MCF7 orthotopic tumors. Early time point (Week 1) and late time point (Week 7) were used to depict non palpable orthotopic tumor stage (small < 2mm) and the palpable tumor stage. Red arrows indicate expected tumor location (mammary gland). Color scales for early lesions (Week 1): 0.2-0.5 SUV-bw; Color scales for large lesions (Week 7): 100-200 SUV-bw. **N.** Relative quantification of radiolabeled 18F-FES uptake in small and large lesions of orthotopic and bone metastases. Each dot represents the mean standard uptake values (mean SUV-bw) of 18F-FES normalized to the mean SUV of 18F-FDG. Mann Whitney *U*-test is used for statistical analysis. n=5 mice per group.

We also tested several other ER+ models in the 3D co-culture assays. MSCs and FOB both induced consistent loss of ER expression across multiple models (Figure 2E-F). In contrast, U937, a human monocytic cell line that is often used to model osteoclast precursors, did not cause the same changes to the ER expression, supporting the specificity of osteogenic cells (Figure 2E).

Hyperactive ER activities can lead to ER protein degradation (Nawaz et al., 1999). Therefore, decreased ER expression could paradoxically suggest an enhanced ER signaling. To examine this possibility, we used real-time qPCR to measure ER at transcription level and found it decreased upon co-culturing with FOB (Figure 2G). Moreover, using a luciferase reporter driven by a promoter containing ER-responsive elements (ERE-luciferase), we discovered that co-culturing with FOB and MSCs indeed decreased ER transcriptional activity (Figure 2H). These data demonstrate that the transient loss of ER is not an indicator of high ER activity, but rather the cause of decreased ER signaling in cancer cells.

Importantly, the MCF7 SCPs also exhibited the same alterations upon interacting with FOB (Figure 2I). In SCP2, SCP3 and SCP4, the degree of ER down-regulation is comparable to parental MCF-7 cells. SCP1, on the other hand, exhibited a lesser decrease (Figure 2J). Interaction with osteogenic cells confers growth advantage on cancer cells as shown in our previous studies (Wang et al., 2015, 2018). SCP1, SCP2, and SCP3 also displayed such advantage in 3D co-cultures as compared to mono-cultures. In contrast, the growth of SCP4 appeared to be suppressed by FOB (Figure S2A). Thus, different SCPs from MCF-7 cells possess variable capacity of orthotopic tumor-initiation, bone colonization, and FOB-mediated growth promotion and ER down-regulation (Figure S2B). This pre-existing heterogeneity supports the importance of clonal selection in metastasis (e.g., in the bone microenvironment, SCP2 and SCP3 are expected to be enriched because of their ability to take advantage of interactions with osteogenic cells), which has been repeatedly demonstrated in previous studies (Bos et al., 2009; Kang et al., 2003; Minn et al., 2005). However, an unappreciated process is the transient changes that occur to most SCPs (Figure 2I-J), independent from the clonal selection.

Positron emission tomography–computed tomography (PET-CT) imaging has been used in clinic to detect bone metastases and evaluate tumor responses to endocrine therapy (Dehdashti et al., 2009). To demonstrate the transient reduction in ER expression in live mice with experimental bone metastases, a radiolabeled 18F-Fluoroestradiol (18F-FES) PET/CT imaging strategy was adopted (Figure 2K). 18F-FES binds ER enriched tumors and can be quantified in parallel with glucose uptake (18F-FDG) to estimate changes in ER expression (Kurland et al., 2017). We found a significant reduction in estrogen uptake in early lesions of bone metastasis (Figure 2L), comparatively to similar lesions in mammary glands (Figure 2M). The discrepancy was reduced in advanced stages of tumors formation –larger tumors (Figure 2N), suggesting a bone specific effect on micrometastatic lesions.

### Interaction with osteogenic cells in the bone microenvironment leads to resistance to endocrine therapies

Downregulation of ER may impact endocrine therapies. To test this hypothesis, we examined the effects of fulvestrant, tamoxifen, and estradiol on ER+ cancer cells with or without co-culture of FOB. The anti-proliferative effects of fulvestrant and tamoxifen, and the proliferative effect of estradiol were significantly blunted in MCF-7 and ZR75-1 cells co-cultures. (Figure 3A and S3A). The presence of FOB also diminished the effects of these agents on ER nuclear localization (Figure 3B).

**Figure 3:**
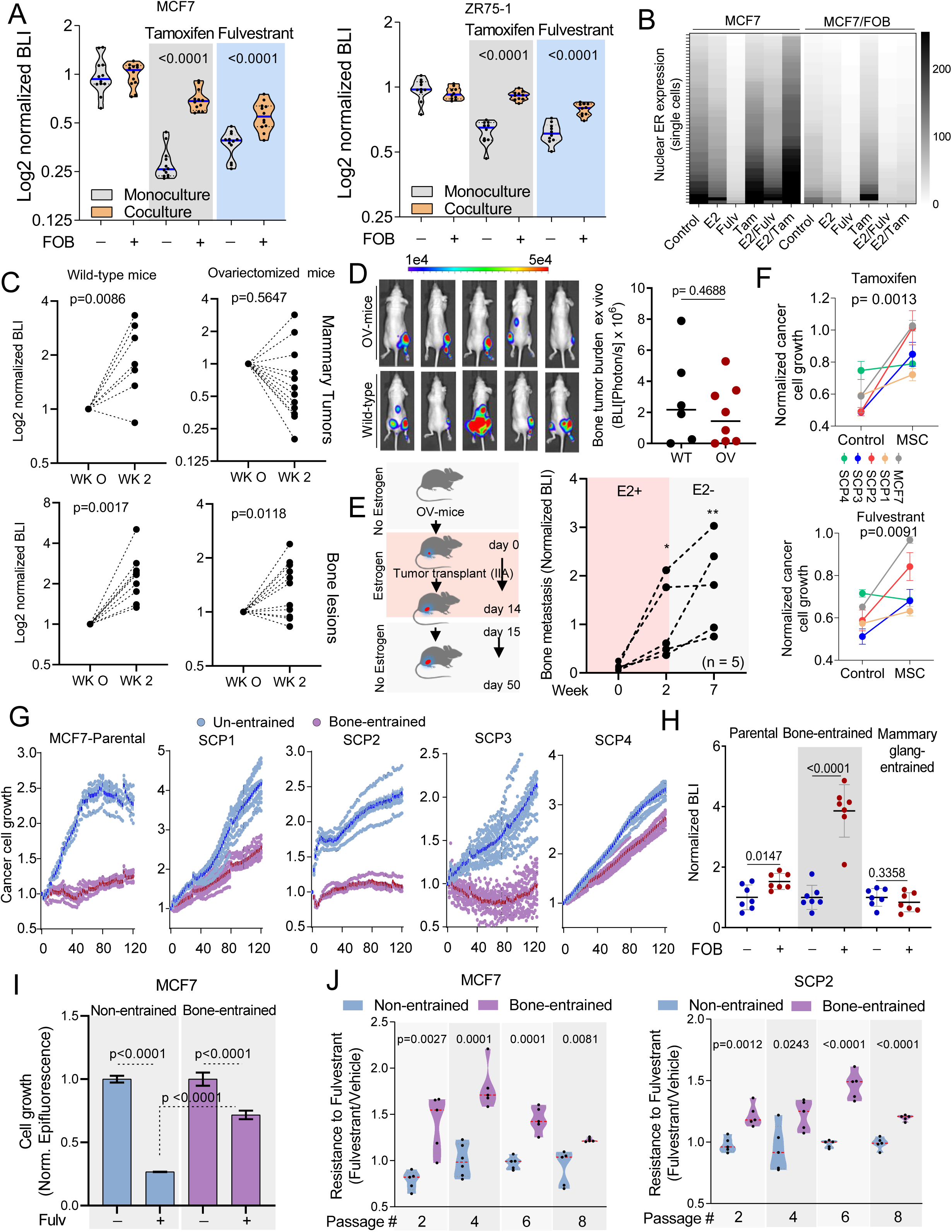
Osteogenic cells confer endocrine resistance **A.** Violin plot showing the response of luciferase-labelled MCF7 and ZR75-1 cells to 100nM of tamoxifen (4-Hydroxytamoxifen) and 20nM of fulvestrant in 3D monoculture or co-culture with osteogenic cells (FOB). Bioluminescence was acquired 72 hours post treatment using the IVIS Lumina II. Two-tailed unpaired Student’s *t*-test was used for statistical analysis (n=12 for MCF7; n=10 for ZR75-1). **B.** Heatmap depicting ER intensity in 3D monocultures and co-cultures of MCF7 cells with FOB following 24 hours treatment with 10nM 17β-estradiol (E2), 20nM fulvestrant (Fulv) and 100nM tamoxifen (4-OHT). Fulvestrant and tamoxifen were used in presence or not of 10nM of E2. Data represent the average of 5 different images. Each row shows the average of 5 random single cells from different images. **C.** Dot plots showing normalized bioluminescence intensity (BLI) of mammary tumor and IIA-induced bone lesions in non-ovariectomized (wild-type) and ovariectomized mice. Ovariectomy was performed 1 month after tumor transplant. Bioluminescence intensity (BLI) was obtained 2 weeks after ovariectomy and normalized to pre-ovariectomy BLI (n= 9 for control groups; n=12 for ovariectomized mammary group; n=11 for ovariectomized bone lesion group). **D.** Bioluminescent images of IIA-induced bone metastases in non-ovariectomized (wild-type) and ovariectomized mice (OV-mice). Different from the experiment described in C, here ovariectomy was performed before IIA-injection of MCF7 cells to mice. 8 weeks after injection, tumor bearing bones were harvested and BLI was assessed ex vivo. **E.** Graphical representation of estrogen replacement via drink water (E2-in-water) for bone metastasis formation in OV-mice. Bone metastasis progression was measured using tumor BLI before and after withdrawal of estrogen from drink water. n=5 mice. P-values ≤ 0.05 and 0.005 (relative to Week 0) are represented as (*) and (**), respectively. **F.** Graphs representing the proliferation of MCF7 (Par) and single cell-derived populations (SCP1-4) in monoculture and MSC co-culture following 1 week of treatment with 20nM fulvestrant and 100nM tamoxifen. n= 5 different cell lines. Two-tailed paired Student’s *t*-test was used for statistical analysis. **G.** Growth kinetics of naïve cells (Parental and SCP1-4) and bone-educated cells (MCF7- Bo, SCP1-Bo, SCP2-Bo, SCP3-Bo, SCP4-Bo) in complete media. n=10 technical replicates. Real-time images were obtained using Incucyte S3 system. **H.** Dot plot representing the BL intensity of bone-entrained (MCF7-Bo), mammary gland-entrained (MCF7-Ma) and naïve MCF7 cells (parental) after 10 days of 3D monoculture (blue) and FOB co-culture (red). Grey area highlight bone-derived (bone-entrained) MCF7 cells. Error bars: mean +/- standard deviation. **I.** Histogram showing the relative growth of non-entrained and bone-entrained MCF7 cells following 20nM fulvestrant treatment. Epifluorescence was measured 72h post treatment. n=12 for bone-entrained, n=4 for non-entrained cells. Two-tailed unpaired Student’s *t*-test was used for all statistical analysis. **J.** Violin plot showing the time course assessment of endocrine resistance phenotype in naïve and bone-entrained MCF7 and SCP2 cells over 8 passages (∼2 months). Median is shown in red. Cells were treated with vehicle or 20nM fulvestrant in estrogen-free medium. n=5-6 technical replicates per group. Two-tailed unpaired Student’s *t*-test was used for all statistical analysis.

We next performed *in vivo* experiments to examine differential responses of ER+ cancer cells to estrogen stimuli in the bone microenvironment. Consistent with previous studies, mammary tumors derived from MCF-7 cells exhibited dependence on estradiol and were stabilized in ovariectomized animals without provision of exogenous estradiol. In contrast, IIA-injected bone lesions of the same cancer cells remained progressive in most ovariectomized animals (Figure 3C). In fact, overiectomy failed to reduce tumor burden or incidence at terminal time point in bones as compared to that in control animals (Figure 3D and S3B). Acute withdrawal of estradiol from animals with established bone lesions did not significantly impact metastatic colonization either (Figure 3E). Interestingly, different SCPs displayed variable resistance to tamoxifen and fulvestrant when interacting with osteogenic cells (Figure 3F). Specifically, whereas SCP2 and SCP3 clearly acquired more resistance to endocrine therapies in co-cultures with FOB, responses of SCP1 and SCP4 did not seem to alter (Figure 3F). This data suggests that the capacity to leverage osteogenic cells and gain resistance to endocrine therapies varies across different clones. However, the resistance itself is not completely pre-existing, but instead induced by osteogenic cells. Therefore, interaction with osteogenic cells induce adaptive alterations in a part of cancer cell clones and render them more resistant to endocrine treatment.

### The bone microenvironment confers “memory”

Since the down-regulation of ER appeared to be reversible (Figure 1), we set out to elucidate the kinetics of this reversal, which is essential for understanding the mechanisms underlying the effects of bone microenvironment. Toward this end, we first established bone lesions using parental MCF-7 cells as well as individual SCPs, and then extracted cancer cells from the bone microenvironment and put them to culture for characterization (termed “bone-entrained” cells). These procedures immediately revealed interesting differences in growth kinetics in vitro – all bone-entrained cells exhibited slower increase over time in culture. This is especially evident for MCF-7 parental, SCP2, and SCP3 cells (Figure 3G). This effect appeared to be bone-specific, as MCF-7 cells extracted from orthotopic tumors or metastatic lesions in other organs exhibited increased (rather than decreased) growth (Figure S3C). When FOB cells are added to the culture, the growth of bone-entrained cells is dramatically increased (Figure 3H and S3D). This newly acquired dependence on osteogenic cells suggests addiction to certain pathways that become activated only in the bone microenvironment.

The bone-entrained cells remained resistance to endocrine therapy in vitro (Figure 3I), and this property last for at least 8 passages for MCF-7 parental cells and SCP2 cells (Figure 3J). Therefore, the reversal of bone microenvironment-conferred effects appear to be relatively slow, suggesting an epigenomic reprogramming process.

### Down-regulation of ER in the bone microenvironment is partially mediated by direct cell-cell contact and gap junctions

We previously reported that heterotypic gap junctions between cancer cells and osteogenic cells mediate calcium influx to the former and activates calcium signaling (Wang et al., 2018). We asked if the gap junction and calcium signaling may mediate ER downregulation and endocrine resistance. This hypothesis was partially validated through western blots showing that suppression of gap junction by a peptide inhibitor, GAP19, or calcium signaling by a small molecule inhibitor, FK506, both partially restore ER expression in co-cultures with FOBs (Figure 4A). This effect was small but noticeable, and was further supported by a converse experiment in which high [Ca2+] in the medium decreased ER expression in MCF7 and ZR75-1 cells (Figure 4B). At the functional level, inhibition of calcium signaling reduced the grow advantage confers by osteogenic cells (FOB) (Figure 4C), and enhanced endocrine therapies in bone-in-culture array (BICA) (Figure 4D), which is an *ex vivo* platform that faithfully recapitulated bone microenvironment and cancer-niche interactions (Wang et al., 2017). Taken together, we provide evidence supporting gap junctions and calcium signaling as one of the mechanisms inhibiting ER expression in bone micrometastases (BMMs).

**Figure 4:**
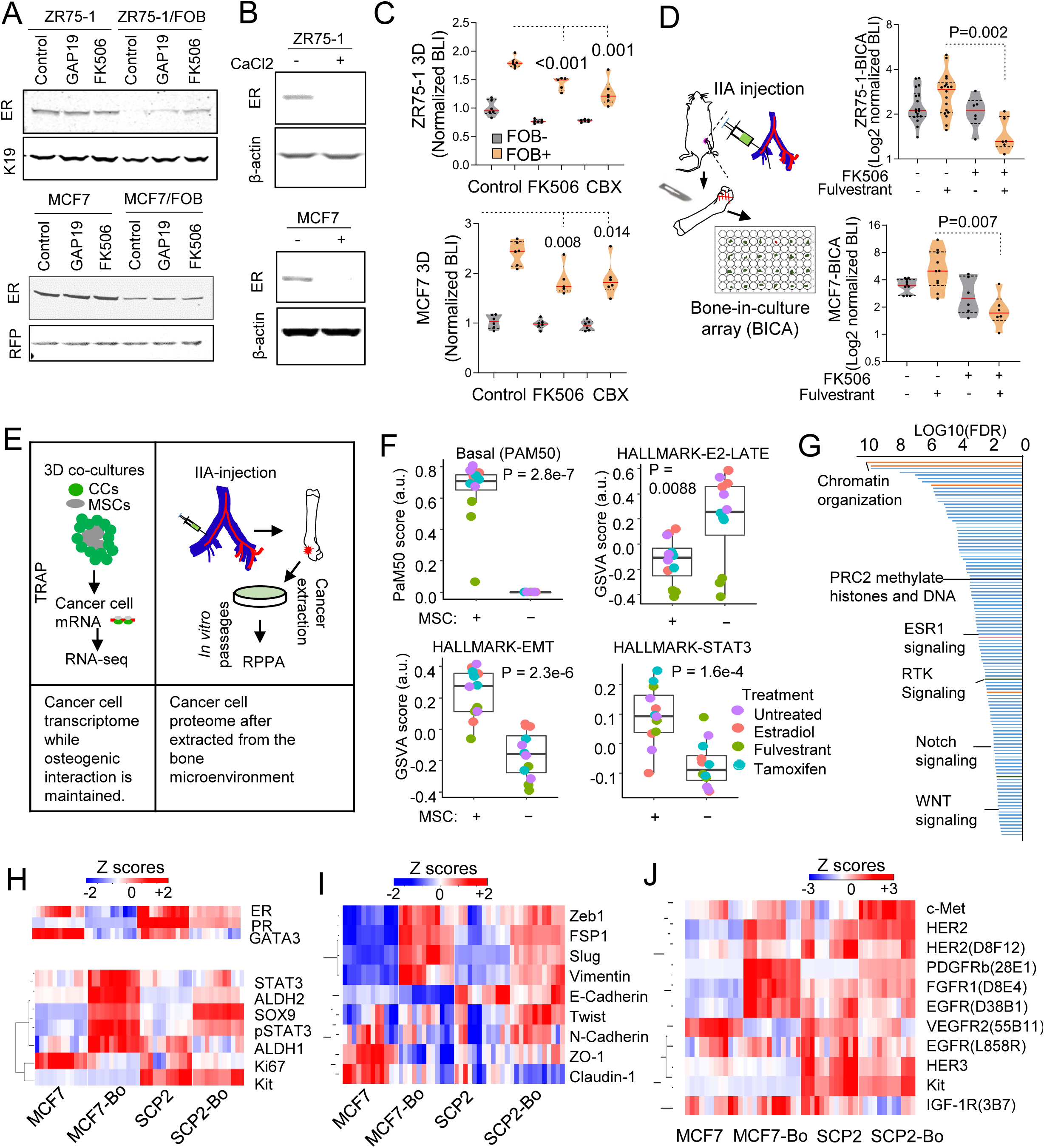
Gap junctions and calcium signaling partially contribute to ER downregulation, and the bone microenvironment drives a global phenotypic shift involving multiple other pathways. **A.** Immunoblotting showing ER expression in ZR75-1 and MCF7 in monoculture or FOB-co-culture after 24h treatment with 1uM CX43 inhibitor (GAP19) and 1uM calcium signaling inhibitor (FK506). Keratin 19 and RFP were used for loading control specific to cancer cells. **B.** Immunoblotting showing the effect of 2mM Calcium (CaCl2) on ER expression in ZR75-1 and MCF7 cells. **C.** Violin plots depicting the inhibitory effect of calcium signaling disruptors (1µM FK506 and 10µM Carbenoxolone-CBX) on osteoblast-induced breast cancer cell growth (ZR75-1 and MCF7). Cancer cells were cultured in 3D with (grey) or without (orange) osteogenic cells (FOB). Bioluminescence intensity (BLI) was acquired 72h post treatment. Data results from 3 different experiments with 4-6 technical replicates. Two-tailed unpaired Student’s *t*-test was used for statistical analysis. **D.** Violin plots indicating synergism between fulvestrant (anti-ER) and FK506 (Calcium signaling inhibitor) on MCF7 and ZR75-1 cells grown in bone using BICA (Bone-In-Culture-Array). Cells were injected to bone using intra-iliac artery (IIA) injection. Hind limbs were harvested, and bone pieces were cultures ex vivo. Bioluminescence intensity (BLI) was assessed using IVIS Lumina II *in vivo* system. n= 6-18 bone pieces for each treatment group. Two-tailed unpaired Student’s *t*-test was used for statistical analysis. **E.** Diagram summarizing strategies used to evaluate molecular changes occurring in cancer cells when exposed to the bone microenvironment. Translating Ribosome Affinity Purification (TRAP) was used to generate and sequence breast cancer cell specific transcriptomes without alteration of cell-cell interaction in 3D co-culture of cancer cells (MCF7) and osteogenic cells (FOB). Reverse Phase Protein Arrays (RPPA) was used to assess protein alterations between naive cells (MCF7 and SCP2) and bone-entrained cells (MCF7-Bo and SCP2-Bo). **F.** Box plot depicting gene signature alternations in MCF7 monoculture (MSC-) and co-cultures (MSC) from TRAP sequencing. Specific colors represent different treatment conditions as indicated. **G.** Waterfall plot showing the gene ontology analysis of TRAP sequencing data PANTHER classification system. Signaling pathways were organized based on their false discovery rate (FDR). **H.** Heatmap depicting expression changes in luminal and stemness-related markers from RPPA data. Parental cells (MCF7 and SCP2), and bone-entrained breast cancer cells (MCF7-Bo and SCP2-Bo) are compared. 4 biological replicates and 3 technical replicates were used for each cell line (See Supplementary Table 1). **I.** Heatmap depicting expression changes in EMT/MET markers from RPPA data as describes in H. **J.** Heatmap depicting expression changes in receptor tyrosine kinases from RPPA data as described in H.

### Unbiased profiling uncovered global phenotypic shift of ER+ cancer cells that persists after dissociation from the bone microenvironment

To identify additional molecular mechanisms underlying ER down-regulation, we used multiple approaches including 1) translating ribosome affinity purification (TRAP) followed by RNA-seq to profile transcriptome in cancer cells that are interacting with osteogenic cells in 3D suspension co-cultures without dissociating the two cell types, and 2) reverse phase protein array (RPPA) to profile over 300 key proteins and phospho-proteins in cancer cells that have been extracted from the bone microenvironment (Figure 4E). In 1), we also applied fulvestrant, tamoxifen and estradiol to the co-cultures to perturb ER signaling. In 2), we included SCP2, a genetically homogenous population that exhibits enhanced ability of bone colonization (Figure S2B) as well as *de novo* bone “addiction” (Figure 3G).

Unbiased hierarchical clustering of TRAP profiling results revealed that in the presence of MSCs the impact of endocrine perturbations became much less evident on ER+ cancer cells as reflected by diminished differences between fulvestrant- and tamoxifen-treated samples and control and estradiol-treated ones (Figure S4A). This supports our previous conclusion that MSCs blunted endocrine responses. We also validated that GJA1, the gene encoding connexin 43, was upregulated by MSCs in co-cultures and exhibited a strong inverse correlation with ER expression (Figure S4B), further indicating a role of gap junctions in down-regulating ER. However, conditioned medium of osteogenic cells also causes ER down-regulation and endocrine resistance (Figure S4C), indicating additional mechanisms based on paracrine signaling.

According to TRAP profiling, over 1,100 genes are significantly increased by MSC co-cultures (FDR < 0.05 and fold change > 2), which is a large number and indicates a global phenotypic alteration. Indeed, using PAM50 signatures, we observed a dramatic shift from luminal to basal subtype (Figure 4F). Consistently, examination of the 50 HALLMARK pathways in MSigDB uncovered several significant changes including the decrease of ER signaling and increase of epithelial to mesenchymal transition (EMT) and STAT3 signaling (Figure 4F), all of which indicated dedifferentiation and stem-like activities(Mani et al., 2008; Marotta et al., 2011; Pfefferle et al., 2015). PANTHER classification system identified a number of pathways overrepresented in the altered genes, including several related to epigenomic regulation of gene expression (e.g., PRC2 activity), stemness-related pathways (e.g., WNT and Notch signaling), and receptor tyrosine kinase (RTK) signaling (Figure 4G, Figure S4D). Some of these pathways have previously been implicated in bone metastasis and therapeutic resistance (Andrade et al., 2017; Esposito et al., 2019; Sethi et al., 2011; Zheng et al., 2017). These findings indicate that the osteogenic microenvironment induces an epigenomic landscape alteration in ER+ breast cancer cells toward more ER-independent and stem-like states.

We used reverse phase protein arrays (RPPA) to molecularly dissect the impact of bone microenvironment that persists even after cancer cells are extracted. We compared the original MCF-7 parental cells and SCP2 with their derivatives that were extracted from bone lesions, which we named “bone-entrained” cells. The proteins and phospho-proteins that are significantly altered were isolated for careful examination (Figure S5A-S5B). The bone-entrained cells, compared to their corresponding controls, exhibited reduced ER signaling or luminal markers (Figure 4H), enhanced stemness (Figure 4H), increased mesenchymal properties (Figure 4I), and strikingly, increased RTK expression (Figure 4J). The most up-regulated protein in both bone-entrained MCF-7 and SCP2 cells are PDGFRβ (Figure S4C). Overall, these indicated a global phenotypic shift toward a more dedifferentiated status (Ginestier et al., 2007; Guo et al., 2012; Mani et al., 2008; Tam et al., 2013; Trastuzumab et al., 2013). Some proteins are expressed at significantly different levels between MCF7 and SCP2, and not altered in bone-entrained cells (e.g., Her3, E-cadherin and Kit), suggesting unique properties of SCP2 which may underlie its enhanced bone colonization capacity. Some other proteins, including PDGFRβ, FGFR1, EGFR, Her2, and c-Met, are significantly upregulated in bone-entrained MCF7 cells. A few of these proteins are already expressed at a higher level in SCP2 (e.g., Her2 and FGFR1), but many exhibited similar elevation in bone-entrained SCP2 (e.g., PDGFRβ, SOX9, pSTAT3, and Zeb1) (Figure 4H-4J). Taken together, the RPPA profiles suggested a mixed action of clonal selection and short-term adaptation during bone colonization. Baseline differences in some proteins suggest that SCP2 is a distinct clone from parental MCF7 cells, and therefore possesses different baseline level of metastasis and *de novo* “addiction” to bone (Figure S2B). However, some others are induced by the bone microenvironment and persist even after leaving the bone microenvironment.

### FGFR and PDGFR pathways contribute to phenotypic changes in BMMs

Among all pathways altered in the bone microenvironment, PDGFRβ and FGFR1 pathways are of particular interest because of their specific implications in breast cancer biology. PDGFRβ exhibited the highest fold change in both models (Figure S5C-S5E) and was shown to determine the subtype of breast cancer and mediate cancer stem cell activities (Lehmann et al., 2011; Tam et al., 2013). Multiple FGF ligands and receptors were found up-regulated in human bone metastases compared to matched primary tumors (Priedigkeit et al., 2017). FGF signaling was also implicated in regulation of stem cell compartment in ER+ breast cancer (Fillmore et al., 2010). This previous knowledge prompted us to further investigate mechanistic links of FGFR and PDGFR signaling to the observed effects induced by the bone microenvironment. Using a literature-based network analysis platform (https://string-db.org/)(von Mering et al., 2005), we found that FGF2 connects ER, FGFR1 and PDGFRB (Figure 5A), suggesting a pivotal role of FGF2 in regulating ER downregulation and endocrine resistance.

**Figure 5:**
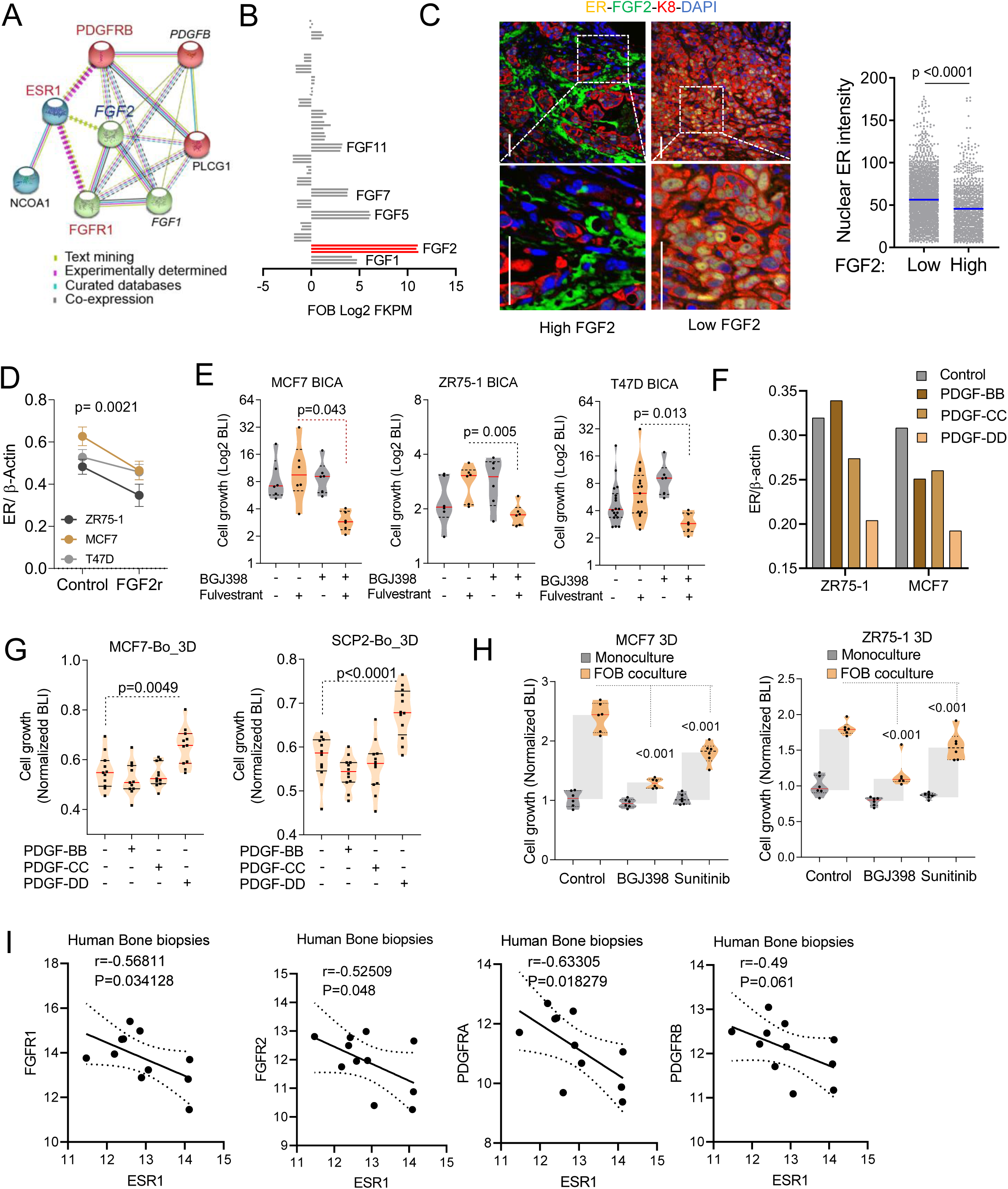
Osteogenic cell-secreted FGFs and PDGFs promote endocrine resistance and bone addiction. **A.** Network depicting functional protein association between FGFR1, PDGFRB and ER using the STRING database. *K*means clustering (*k*=3) was used to represent 3 major centroids (depicted as red, green, and cyan spheres) and their most closely associated proteins based on unsupervised data mining. **B.** Graph showing the gene expression of human FGFs in osteogenic cells (FOB). The data derives from RNA sequencing data of osteogenic cells (FOB). n= 3 technical replicates **C.** Representative IF images showing decreased ER expression (yellow) in tumors established in FGF2 (green) enriched bone microenvironments. Keratin 8 (red) is used to identify breast cancer cells. The scatter dot plot represents ER quantification from tumors according to FGF2 enrichment (Low and High) in adjacent stromal cells (n=3-4 samples). Mean expression is represented in blue. **D.** Graph depicting the inhibitory effect of recombinant FGF2 (20ng/ml) on ER expression in MCF7, ZR75-1 and T47D after 24h treatment. Data represent normalized ER over β-actin protein expression from 3 separate immunoblotting experiments. **E.** Bone-In-Culture-Array (BICA) assay showing synergistic effects between 2.5µM FGF2 inhibitor (BGJ398) and 20nM fulvestrant in MCF7, ZR75-1 and T47D models. **F.** Graph depicts the inhibitory effect of indicated PDGF recombinants on ER expression in MCF7 and ZR75-1 after 24h treatment. Data represent normalized ER over β-actin protein expression from Immunoblot quantification. **G.** Violin plot showing the effect of 20ng/ml PDGF recombinants (PDGF-BB, PDGF-CC, PDGF-DD) on bone-entrained MCF7 (MCF7-Bo) and SCP2 (SCP2-Bo) response to fulvestrant. BLI was assessed after 72h of treatment. **H.** Dot plot showing the effect of the pan FGFR inhibitor (BGJ398), and the PDGFRβ inhibitor (sunitinib) on osteoblast (FOB)- mediated MCF7 and ZR75-1 cell growth in 3D. **I.** Scatter plot showing negative Pearson correlation (r) between FGFR1, FGFR2, PDGFRA, PDGFRB, and ESR1 expression in clinical specimens of bone metastasis. n= 11 bone metastasis samples. *P* values: two tailed Paired Sample *t* Test.

Indeed, FGF2 is the highest expressed FGF ligands by FOB cells (Figure 5B). Immunofluorescence staining of FGF2 on bone specimens revealed an inverse correlation with nuclear intensity of ER in BMMs (Figure 5C). Functionally, recombinant FGF2 treatment decreased ER expression in multiple cell lines (Figure 5D), whereas a potent FGFR inhibitor, BGJ398, reversed fulvestrant resistance of ER+ cancer cells in BICA (Figure 5E).

On the other hand, PDGFR signaling also appeared to regulate ER expression. Specifically, PDGF-DD, but not PDGF-BB and PDGF-CC, seemed to significantly down-regulate ER expression (Figure 5F), consistent with the specific upregulation of PDGFRB in bone-entrained cells. Further experiments uncovered a potent effect of PDGF-DD on cell growth of bone-entrained MCF-7 and SCP2 cells (Figure 5G), which might partially explain the “addictiveness” of these cells to the bone microenvironment (Figure 3H), a PDGF-enriched milieu. Like inhibition of FGFR, inhibition of PDGFR signaling by sunitinib also partially abolished the promoting effects of FOB cells on cancer cell growth in 3D co-cultures (Figure 5H), further supporting the important roles of both FGFR and PDGFR signaling in the interaction between cancer cells and the osteogenic niche.

In clinical bone metastases, we observed significant inverse correlations between ESR1 expression and multiple FGFR and PDGFR genes (Figure 5I). Taken together, these data strongly suggest that transient ER downregulation and phenotypic shift of cancer cells in the bone microenvironment may be driven by paracrine RTK signaling, especially FGFR and PDGFR.

### The complicated impact of bone microenvironment converges on an EZH2-mediated phenotypic-shift of ER+ breast cancer cells

We next asked how the discovered pathways cooperate to influence the epigenomic landscape of ER+ bone micrometastases, and in turn silence ER and cause a luminal-to-basal phenotypic shift. Using the Epigenomic Roadmap database, we discovered that FGF2-regulated genes are predominantly enriched with tri-methylation of H3K27, and sensitive to perturbation of EZH2 (Figure 6A). This is consistent with the finding that PRC2 methyltransferase activity is enhanced in cancer cells co-cultured with MSCs (Figure 4G). Indeed, treatment of both recombinant FGF2 and PDGF-DD increased H3K27me3 and EZH2 expression in SCP2 cells (Figure 6B), but do not significantly affect other H3 modifications (Figure S6A). Conversely, treatment of BGJ298 decreased EZH2 expression in 3D cancer-MSC co-cultures (Figure 6C). Furthermore, it appeared that calcium signaling may also affect EZH2 expression at the RNA level (Figure S6B). Thus, the pathways that were discovered to downregulate ER seem to converge on the regulation of EZH2.

**Figure 6:**
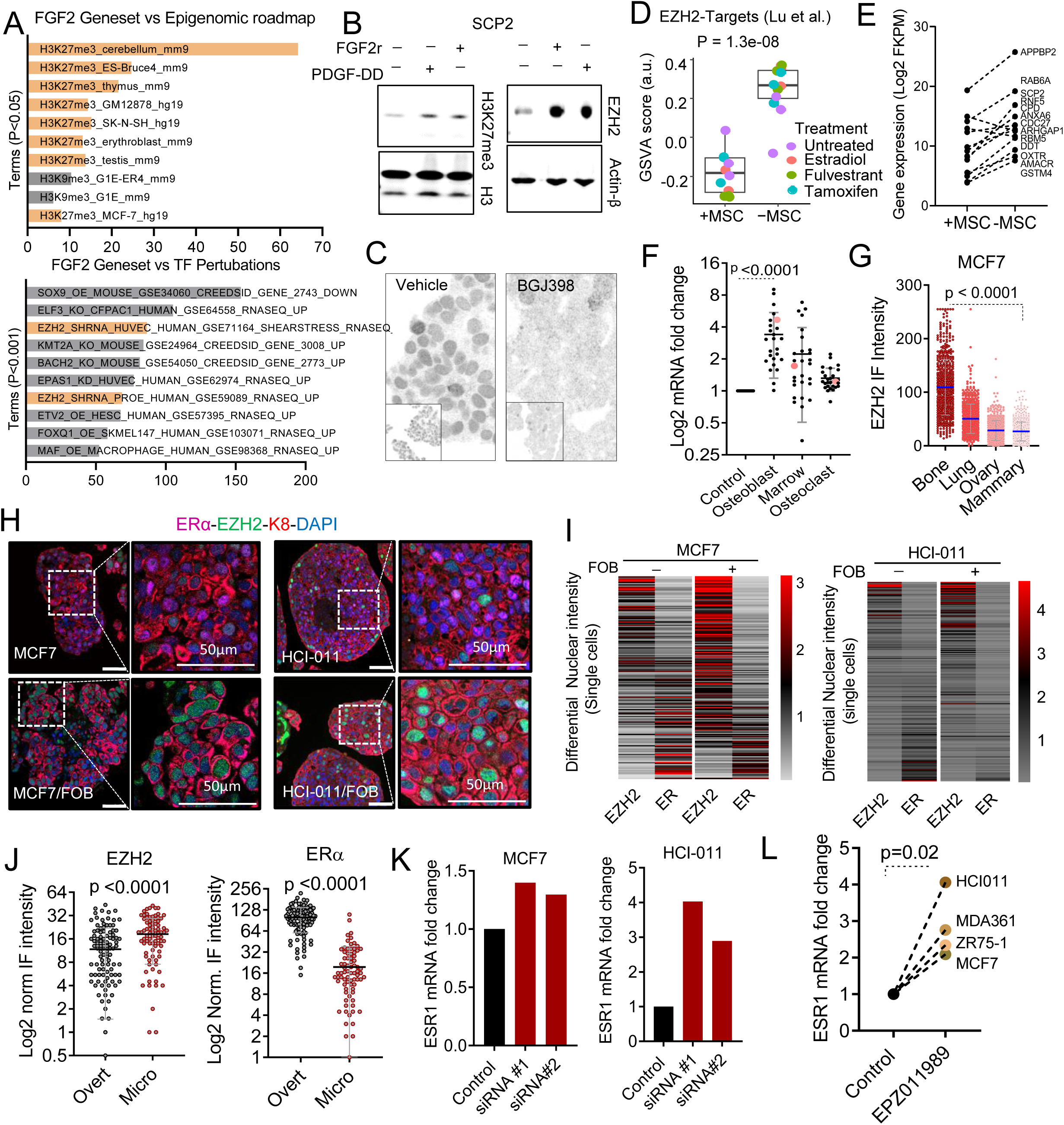
EZH2 integrates multiple signals from the bone microenvironment and drives the phenotypic shift of ER+ breast cancer cells. **A.** Graphs showing the association of histone modifications with FGF2 gene signatures using the Enrichr platform (https://amp.pharm.mssm.edu/Enrichr/). Processed ChIP-sequencing data was obtained from epigenomic roadmap project (Roadmap Epigenomics Consortium et al., 2015). Signatures are sorted based on p-value ranking. Only p-value < 0.05 and <0.01 were shown for the top and bottom panel, respectively. **B.** Immunoblotting showing changes in H3K27me3 and EZH2 expression in nuclear extracts of SCP2 cells following 24h treatment with 20nM FGF2 recombinant (FGF2r). Total histone 3 (H3) and actin-β were used as loading controls for H3k27me3 and EZH2, respectively. **C.** Primary cells generated from HCI011 (ER+ PDX) were cultured in 3D and treated with 1µM pan FGFR inhibitor (BGJ398) or vehicle for 24h. Representative images depict EZH2 expression in indicated conditions. **D.** Box plot representing the gene set variation score (GSVA) of EZH2 target genes (Lu et al.) in MCF7 monoculture (MSC-) and co-culture with MSCs (MSC+) from TRAP-sequencing. Each color represents a specific treatment as indicated. Cells were cultured (estrogen-free medium) in 3D and treated with 10nM estradiol and fulvestrant, and 100nM tamoxifen for 24h. **E.** Graph shows reductions in EZH2 target gene expression in MCF7 cells following 3D monoculture (-MSC) and co-culture (+MSC) and TRAP sequencing. EZH2 signature genes were selected from previous studies (Zhou et al., 2002). **F.** Quantitative PCR of stemness-related genes in MCF7 cells from 3D monoculture and co-culture with FOB (osteoblast), human bone marrow (marrow), and U937 (osteoclast) cells. All conditions were FACS-sorted for RFP-labeled MCF7cells before mRNA extraction and qPCR. **G.** IF quantification of EZH2 expression in multiple metastases and primary tumor originating from the same mouse. 2×10^5^ MCF7 cells were transplanted orthotopically and via IIA injection (bone) to nude mice, which led to tumor formation at multiples sites (Lung, ovary, bone and Mammary gland). Metastases derived from same animal were harvested for immunofluorescence studies. **H.** Representative confocal images showing co-expression of ER (purple) and EZH2 (Green) in 3D models of MCF7 and HCI011 primary cells. Keratin 8 (red) and DAPI (Blue) were used to identify epithelial cells and cell nuclei. **I.** IF quantification of ER (purple) and EZH2 (green) in MCF7 and HCI011 3D monocultures and co-cultures with osteogenic cells (FOB). Keratin 8 -K8 (red) and DAPI (blue) were used to identify epithelial cells and cell nuclei. **J.** IF quantification showing changes in EZH2 and ER according to bone metastasis size. Micrometastases (micro) represent early stages while macrometastases (overt) represent late stages of bone metastasis. **K.** Quantitative PCR showing the effect of siRNA downregulation of EZH2 on ESR1 expression. n= 3 technical replicates. **L.** Quantitative PCR showing the effect of EZH2 inhibitor EPZ011989 on ESR1 expression after 24 hours of treatment. n=4 cell lines. P value shows two-tailed paired Student’s *t* test.

The downstream PRC2 target genes are concertedly downregulated by co-culturing of MSCs (Figure 6D-6E). EZH2 is a reliable marker for cancer stemness (Kim and Roberts, 2016; Malta et al., 2018). Consistently, a stemness signature exhibited markedly increased expression in co-cultures with osteoblasts and bone marrow cell including MSCs, but not with osteoclasts (Figure 6F). In addition, EZH2 expression appeared to be specifically enhanced in cancer cells residing in the bone microenvironment as compared to the same cancer cells in other organs (Figure 6G). Together, these data confirmed enhanced EZH2 activities in the bone microenvironment, which are mediated by interaction with osteogenic cells.

To ask if EZH2 silences ER expression and causes the luminal-to-basal phenotypic shift, we carried out IF staining in 3D co-cultures and revealed that interaction with FOB induced EZH2 expression (Figure 6H). Moreover, at a single cell level, EZH2 expression inversely correlates with ER expression both in 3D cultures and in bone lesions (Figure 6I and 6J). Knockdown of EZH2 by siRNAs or inhibition of EZH2 enzymatic activity by EZH2 inhibitor (EPZ011989) (Campbell et al., 2015) led to restoration of ER expression at the RNA level (Figure 6K, 6L, and S6C). In human datasets, the expression of EZH2 target genes correlates with distant metastasis-free survival (DMFS) in ER+ and tamoxifen-treated patients, but not in ER-negative (Figure S6D), and inversely correlates with ESR1, as well as canonical ER target genes including PR, GATA3, TFF1 and FOXA1 (Figure S6E). Collectively, these data identified EZH2 as a key regulator that integrate multiple upstream signals from the bone microenvironment and in turn drive the phenotypic shift of metastatic cells.

### Short-term inhibition of EZH2 restores sensitivity of bone micrometastases to endocrine therapies

Since EZH2 mediates bone microenvironment-induced endocrine resistance, we hypothesize that inhibition of EZH2 should reverse this resistance and synergize with endocrine therapies. This is indeed the case in vitro. The synergy is especially strong on bone-entrained MCF-7 cells (Figure 7A). We also conducted *in vivo* experiments to test this hypothesis. A four-arm experiment was used to specifically ask if combinatory treatment of EPZ011989 and fulvestrant at the microscopic metastasis stage (to mimic adjuvant therapy) could lead to decreased bone colonization (Figure S7A). At overt metastasis phase (4 weeks after “adjuvant” treatment), fulvestrant was again used for treatment to control tumor progression. In this setting, EPZ011989 and fulvestrant had little or modest effects respectively as single agents. However, the combined treatment strongly inhibited bone colonization and rendered 50% of animals tumor-free by the end of experiment (Figure 7B-D). This is a remarkable effect considering that EPZ011989 treatment only last for 3 weeks.

**Figure 7:**
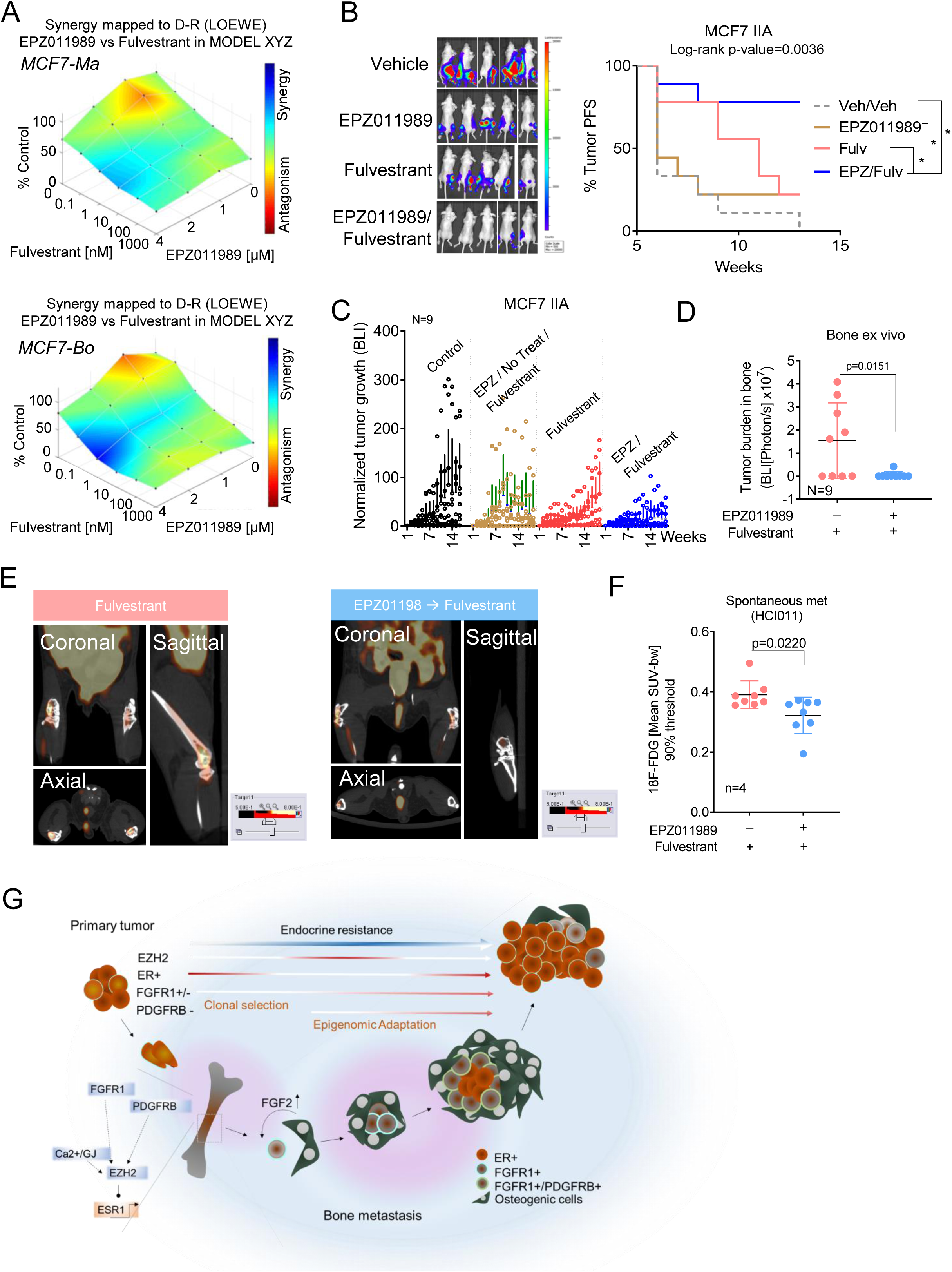
Short-term treatment of an EZH2 inhibitor restores endocrine sensitivity of breast cancer bone metastasis. A. LOEWE analysis of fulvestrant and EZH2 inhibitor (EPZ011989) combination in 3D co-culture of mammary gland-entrained (MCF7-Ma) and bone-entrained (MCF7-Bo) MCF7 cells. Graphs were generated using the Combenefit interactive platform(Di Veroli et al., 2016). B. Representative images showing beneficial effects of fulvestrant and EPZ011989 combination treatment on IIA-induced bone metastases. 4 arms were formed: (i) control group (Ctrl/ctrl) received vehicle treatment; (ii) EPZ011989 group (EPZ011989) received EZH2 inhibitor followed by vehicle treatment; (iii) fulvestrant group (Fulv) received vehicle treatment followed by fulvestrant treatment; (iv) combination treatment group (EPZ/Fulv) received both EPZ011989 and Fulvestrant treatment. EPZ011989 was used as a neoadjuvant for 3 weeks (125mg/kg; oral gavage; twice a day) before fulvestrant treatment (250mg/kg; subcutaneous injection, once per week for 2 weeks). Tumor progression free survival (PFS) curve was generated to represent the effect of indicated treatments. Log-rank (Mantel-Cox) test was used for statistical analysis. P values <0.05 are represented as (*). C. Graph showing the bioluminescence intensity of IIA-induced bone metastases normalized to day 1 of treatment. Each dot represents an image. D. Ex vivo quantification of tumor bioluminescence intensity in bone. E. Representative PET-CT images showing 18F-FDG uptake in hind limb bones of fulvestrant treated and EPZ011989/fulvestrant treated metastases. 2×10^5^ dissociated tumors cells from freshly harvested PDXs (HCI011) were injected to mammary gland of nude mice. A week after primary tumors were removed, EPZ011989 treatment (125mg/kg; oral gavage; twice a day) started for 3 weeks, followed by fulvestrant treatment (250mg/kg; subcutaneous injection, once per week) for 2 weeks. Residual tumors were challenged with estrogen supplementation in drink water before 18F-FDG PET-CT imaging. F. Quantification of 18F-FDG uptake (mean SUV-bw) in hind limbs to evaluate spontaneous metastasis from HCI011 PDXs. A 90% thresholding of the maximum standard uptake value (SUV-bw) was used to remove background signals. Two-tailed unpaired Student’s *t*-test was used for statistical analysis. G. Proposed model summarizing the mechanisms involved in breast cancer bone metastasis and endocrine resistance. A subset of ER+ breast cancers express FGFR1 which may confer survival advantages during metastasis. FGFs are important for stem cell maintenance and migration. Here, we found that FGF2 secreted from osteoblasts can activate FGFR signaling in a paracrine manner, leading to increased EZH2 expression, chromatin alteration, and subsequent downregulation of estrogen receptor (ER) in the osteogenic niche. This promotes estrogen independent growth of micrometastases. The proximity of cancer cells to osteoblasts is gradually lost with macrometastasis formation resulting in ER re-expression in advanced tumors. These changes are also associated with acquired PDGFRB expression which further contributes in maintaining stemness in overt lesions of bone metastases. As PDGFRB expression persists outside of the bone microenvironment, it could be a potential marker for bone metastasis.

We also tested the combinatory treatment on PDX-based spontaneous bone metastasis models using PET imaging. Pretreatment of mice with EPZ011989 inhibited spontaneous metastasis to bone as shown by the reduced 18F-FDG update (Figure 7E-7F). Similarly, 18F-NAF distribution in skeletal bone revealed less bone metastasis (Figure S7C).

## DISCUSSION

Phenotypic plasticity has been increasingly recognized as a major driving force of normal development, tumor initiation, and tumor progression (Dravis et al., 2018; Gupta et al., 2019; Lambert et al., 2017). In this study, we have uncovered that the osteogenic cells trigger a global epigenomic change in ER+ metastatic seeds through both paracrine signaling and direct cell-cell contact. Importantly, this global change represents an adaption to the bone microenvironment, and leads to increased phenotypic plasticity and therapeutic resistance.

These changes form a stable “memory” that persists even after cancer cells are removed from bone and grown *in vitro*. The fact this “memory” occurs to genetically homogeneous populations distinguishes it from clonal selection, which has been intensively investigated in the past (Bos et al., 2009; Kang et al., 2003; Minn et al., 2005b). Indeed, our data support a coordinated action between epigenomic adaptation and genetic selection. Specifically, genetic traits (e.g., expression of FGFR1) may determine the capacity of a cancer cell to undergo further epigenomic alteration (e.g., up-regulation of PDGFR and EZH2). Our findings were supported by ER+ PDX models of bone metastasis, which remains unprecedented to our knowledge.

Our study identified a number of pathways that are altered in cancer cells by the bone microenvironment. Among these pathways, EZH2-mediated epigenomic reprogramming is a leading candidate for therapeutic intervention. It integrates multiple signals from osteogenic cells (e.g., FGF2 and PDGF-DD), and in turn, broadly impacts several downstream pathways related to cancer stemness and metastasis (e.g., WNT and Notch)(Gonzalez et al., 2014; Shi et al., 2007). Moreover, potent and selective EZH2 inhibitors are available and being clinically investigated in other diseases (Italiano et al., 2018), making it relatively easy for future clinical applications. Pharmacological inhibition of EZH2 promotes a global landscape change of histone marks (Huang et al., 2018). Tumors developed resistance to histone demethylase KDM5A/B had increased EZH2 expression (Hinohara et al., 2018). Hence, the bone microenvironment induction of EZH2 in BMMs may trigger an epigenomic disturbance beyond H3K27me3.

The loss of ER expression during bone metastasis appears to be transient. In the advanced stage when the osteolytic vicious cycle starts (Boyce et al., 1999; Kozlow and Guise, 2005; Weilbaecher et al., 2011), ER expression seems to recover, which might be caused by the opposite effects of several other cell types (e.g., represented by U937 and Hs5 in Figure 2E) that are recruited to metastases later. However, the microenvironment-conferred endocrine resistance persists for several cell cycles even after cancer cells are dissociated from the bone milieu. Thus, overt bone metastases may be heterogeneous, including a subset whose ER signaling remains repressed, which may be responsible for rapid reappearance of resistance. This possibility is supported by our initial observation in spontaneous bone metastasis from PDX tumors (Figure S1A). Alternatively, endocrine resistance may be driven by additional ER-independent mechanisms, and therefore, recovery of ER cannot fully restore sensitivity. In either case, overt bone metastases may be ER+ and partially sensitive to endocrine therapies, but resistance can quickly develop – a phenomenon mimicking the clinical reality (Johnston, 2010).

Unlike FGFR1 which is expressed at a higher level in some SCPs before reaching the bone, PDGFRB expression appears to be activated by the bone microenvironment. Functionally, PDGFRB was shown to mediate stem cell-specific signaling and drive stroma-induced subtype-shift (Roswall et al., 2018; Tam et al., 2013). Here, our data suggest that it contributes to phenotypic plasticity and endocrine resistance. It might also be used as a cell surface marker of cancer cells ever lodged to the bone. Similar approaches may be applied to identify markers of epigenetic imprints on metastatic cancer cells in other organs. Ultimately, this information may allow us to predict the location of metastasis by examining these imprints on CTCs.

Although our experiments focused on bone metastases, we are not ignoring the fact that other metastases also need to be prevented and cured. In fact, patients with bone-only metastases have better prognosis than patients with metastases in other more vital organs (Coleman, 2001). However, bone may not be the final destination of dissemination. Recent genomic analyses revealed frequent metastasis-to-metastasis seeding (Brown et al., 2017; Ullah et al., 2018). Over two-thirds of bone-only metastases subsequently develop other metastases (Coleman, 2001). The finding that stem cell signaling is elevated in the bone microenvironment actually raises the possibility that bone may invigorate disseminated tumor cells for further metastases, and this possibility has recently gained support in our co-submitted manuscript (Zhang et al., 2019). Therefore, investigations on bone metastasis may have broader impact.

## Acknowledgements

We thank all members of the Zhang Lab for their insightful contributions. Our thanks also go to the Breast Center Pathology Core, the RPPA core, the Small Animal Imaging Facility (SAIF) core for their technical support, and to all members of Lester and Sue Smith Breast Center. We express our gratitude to Dr. Rosen Jeffrey for his invaluable assistance and guidance, to Drs. Michael Lewis, Keith Chen, Fotis Nikolo, Xi Chen, Jin Cao, Fabio Stossi, Michael Mancini, and to Lacey Dobrolecki for their technical support. We finally thank Epizyme (Cambridge, MA) for kindly providing the EZH2 inhibitor EPZ-011989.

## Author contributions

Conceptualization, I.B., G.M.W., H.S, H.W., P.S, and X.H.-F.Z.; Methodology: I.B., H.W., P.S., W.Z., J.L., W.U., H.C.L, A.M., M.J., I.K., S.S., A.G., P.Si., H.S., G.M.W., M.J.E., X.H-F.Z; Software and Data Curation, I.B., H.W. and X.H-F.Z. Resources, M.J.E, and X.H.-F.Z.; Writing-original Draft: I.B., and X.H.-F.Z.; Writing – Review & Editing, I.B., H.W., W.Z., and X.H.-F.Z.; Visualization, I.B. and X.H.-F.Z.; Project Administration, I.B., J.L. and X.H.-F.Z.; Supervision and Funding Acquisition, X.H.-F.Z.

## List of Supplementary Figures and Legends

**Supplementary Figure 1, related to Figure 1:**
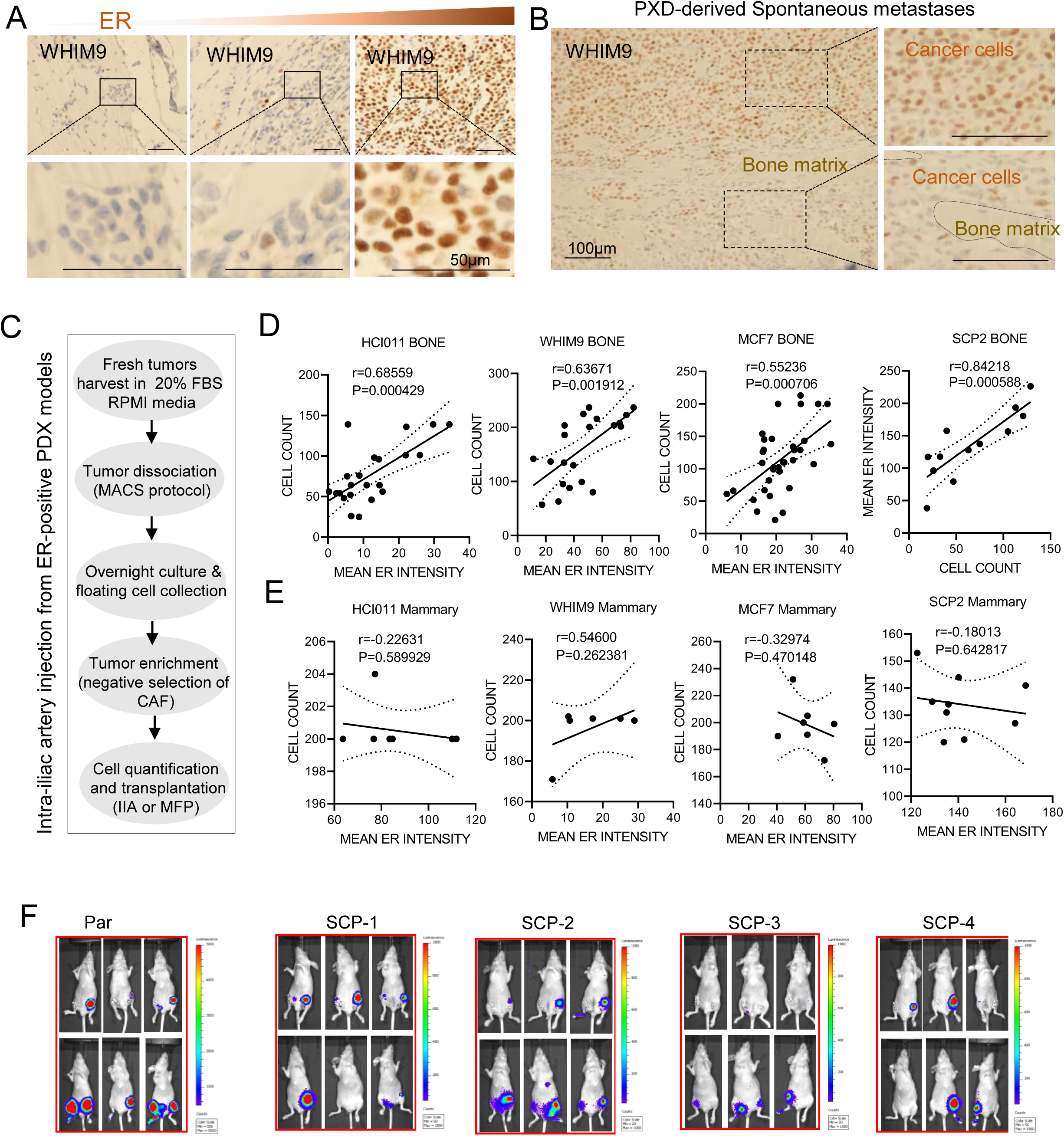
A. ER IHC staining are shown for spontaneous metastasis of WHIM9, respectively. Representative pictures of metastatic lesions of different sizes are shown. Scale bar: 50μm. B. ER IHC staining of a macroscopic spontaneous metastasis is shown with bone matrix annotated. Scale bar, 100μm. C. Diagram representing key procedures for intra-iliac artery injection (to generate bone metastasis) from freshly harvested orthotopic PDX models. D. Scatter plots showing Pearson correlation (r) between bone metastasis sizes (cell count) and ER IF intensity. Images were acquired with 40x oil objective lens (Leica TCS SP5 confocal microscope). Each dot represents an image (n=22 for HCI011; n=21 for WHIM9; n=34 for MCF7; n=12 for SCP2) E. Scatter plots showing Pearson correlations (r) between orthotopic tumor sizes (cell count) and ER IF intensity. Images were acquired with 40x oil objective lens (Leica TCS SP5 confocal microscope). Each dot represents an image (n=8 for HCI011; n=6 for WHIM9; n= 7 For MCF7; n=9 for SCP2). *P* values: two tailed Paired Sample *t* Test. F. Representative images of orthotopic and IIA-induces bone metastasis from MCF7 and MCF7 single cell-derived population (SCP1-4). From the ventral view, right tumors show orthotopic tumors while left tumors are bone metastases.

**Supplementary Figure 2, related to Figure 2:**
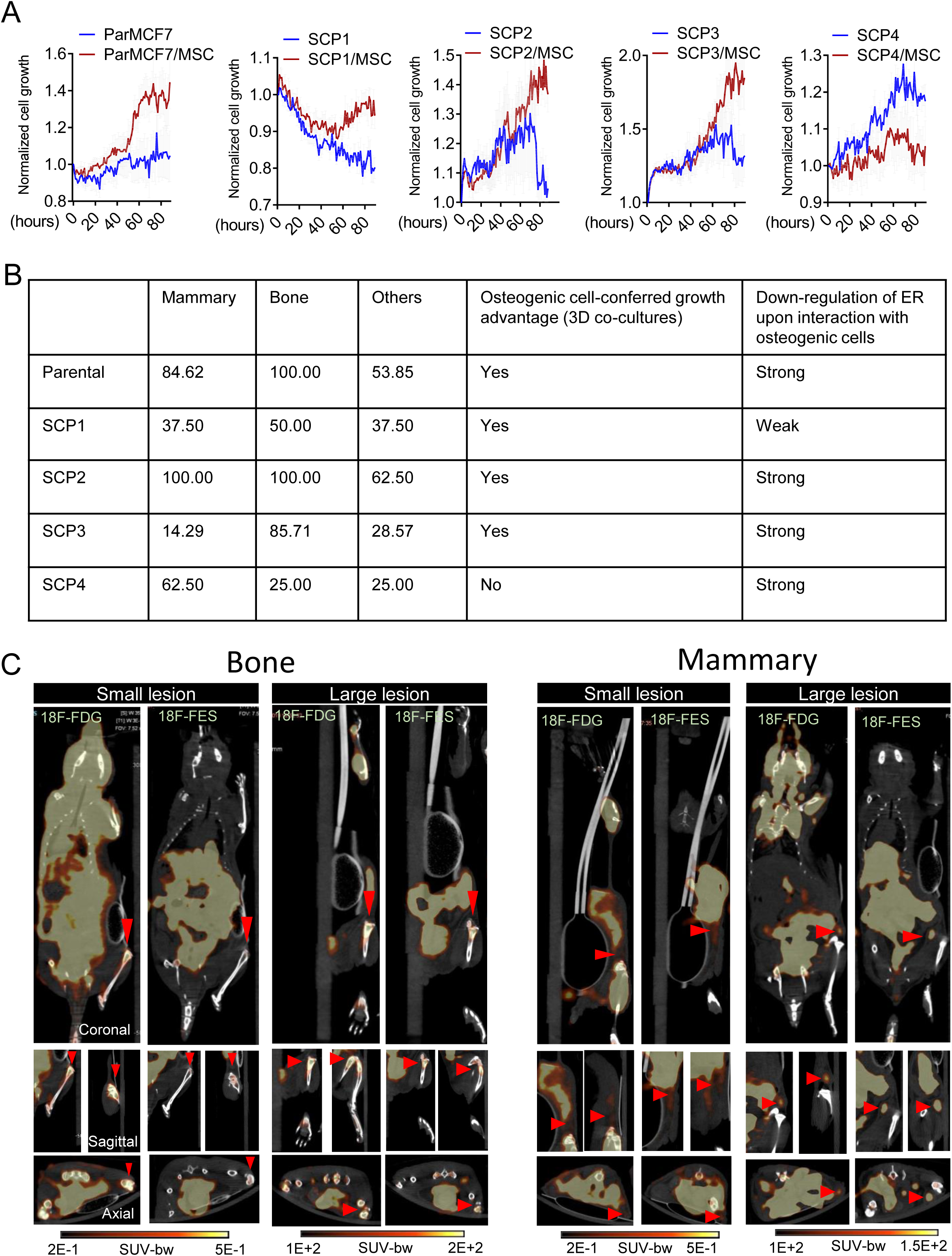
A. Growth curves of MCF7 and single cell-derived populations (SCP1-4) in 3D monoculture and co-culture with osteoprogenitor cells (MSC) in complete growth medium (10% FBS). Real-time images were acquired hourly using Incucyte. Values represent epifluorescence normalized to the earliest time point of co-culture. Error bars: +/- standard error of the mean (SEM). B. Table summarizing metastatic characteristics of MCF7 (Parental) and SCPs in vivo and in 3D co-culture C. Coronal, sagittal, and axial view or representative PET/CT images depicting the uptake of radiolabeled fluorodeoxyglucose (18F-FDG) and fluoroestradiol (18F-FES) in small and large lesions of MCF7 orthotopic and bone metastasis. Small lesions (Week 1) and larger lesions (Week 7) were used to depict early and late stage of tumor formation. Red arrows indicate expected tumor location in bone and mammary gland. Color scales for early lesions (Week 1): 0.2-0.5 SUV-bw; Color scales for large lesions (Week 7): 100-200 SUV-bw.

**Supplementary Figure 3, related to Figure 3:**
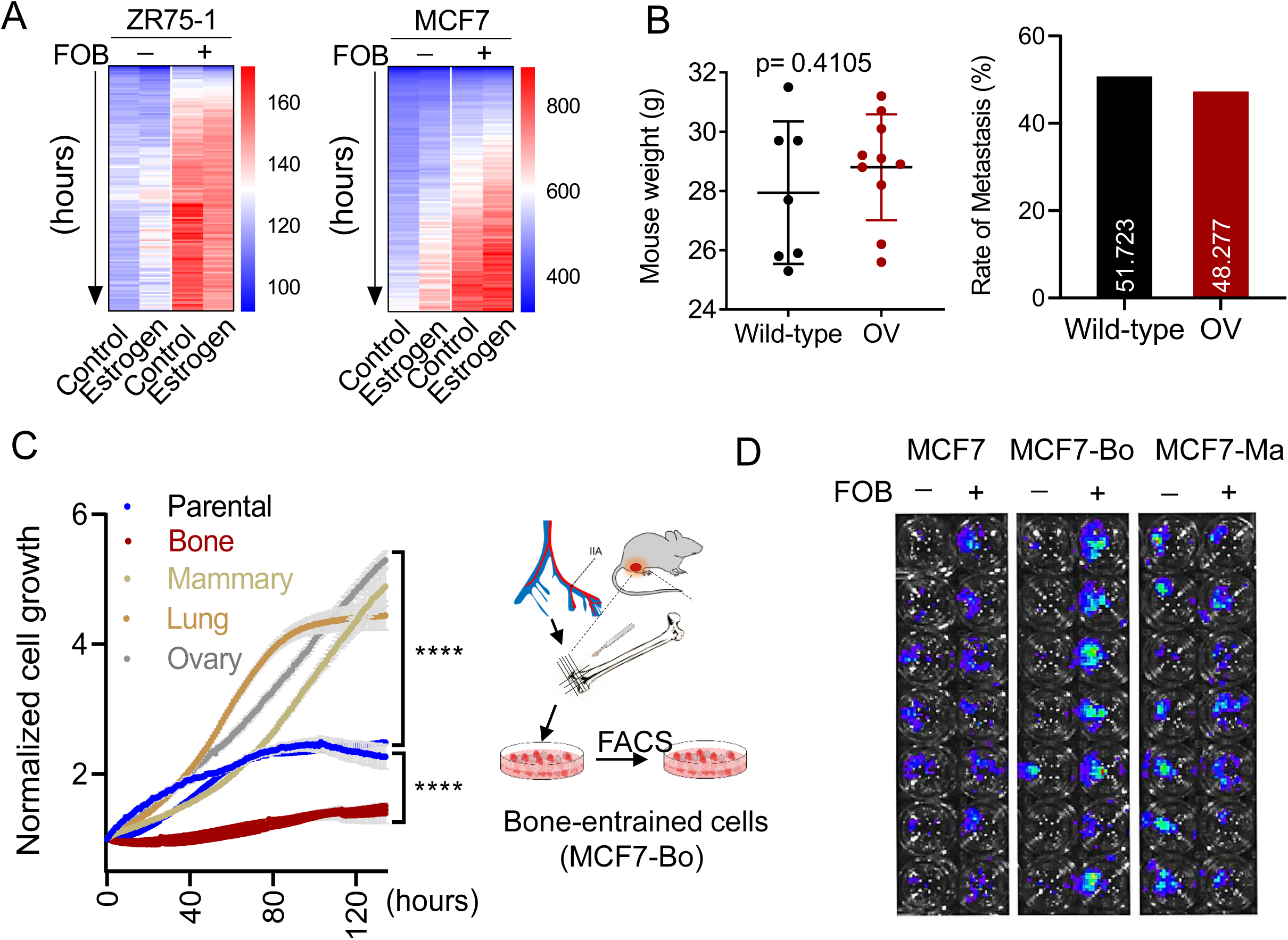
A. Heatmap showing estrogen independent promoting effect of osteogenic cells (FOB) on cancer cells (MCF7) in 3D co-culture. Real-time epifluorescence images were acquired with Incucyte live imaging for 6 days. n=3-5 replicates per condition. P values were calculated using ordinary two-way ANOVA (p<0.0001 for ZR75-1 and MCF7). B. Dot plot showing the effect of ovary removal (OV) on mouse weight in comparison to wild-type mice (Left), and the effect of OV on the rate of bone metastasis formation (right). For wild-type group, n= 8 mice; for OV group n= 6 mice). *P* values: two tailed Paired Sample *t* Test (p-value <0.05 is significant). C. Graph showing growth differences in MCF7 cells derived from different tissues. The diagram represents the work flow used to generate bone-entrained cells (MCF7-Bo). D. Representative luciferase images showing stimulatory effect of FOB on non-entrained (MCF7), bone-entrained (MCF7-Bo), and mammary gland-entrained (MCF7-Ma) cells. Images are acquired at day 10 post co-coculture.

**Supplementary Figure 4, related to Figure 4:**
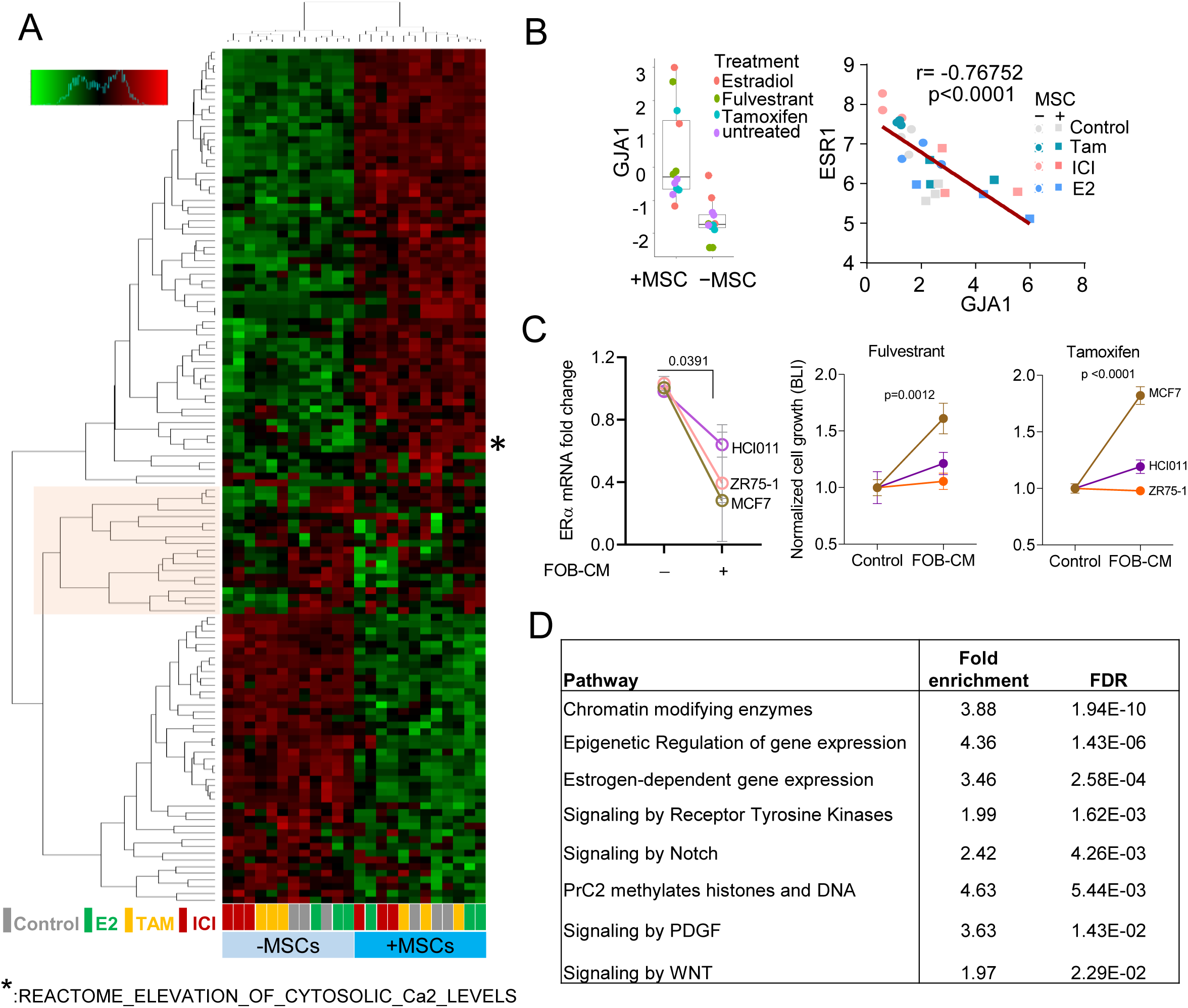
A. Heatmap depicting molecular pathways (PANTHERS) altered in MCF7 when cultured alone or in 3D with MSC and treated with vehicle (control), 10nM 17B-estradiol (E2), 10nM fulvestrant (ICI) or 100nM tamoxifen (Tam) for 24h. Data results from Translating Ribosome Affinity Purification (TRAP) sequencing analysis. B. Boxplot showing the promoting effect of osteogenic cells (MSC) on CX43 gene (GJA1) expression in 3D co-cultures of MCF7 (left). Scatter plot showing Pearson (r) correlation between GJA1 and ESR1 gene (right). All values result from Translating Ribosome Affinity Purification (TRAP) sequencing data. Colors are specific to treatment conditions. C. Effect of FOB conditioned media on ESR1 expression (left) and endocrine response. n=3 different cell models. For left panel (mRNA expression), *P* values: two tailed Paired Sample *t* Test (p-value <0.05 is significant). For cell growth measured by BLI (right panel), *P* values were calculated using ordinary two-way ANOVA. D. Table depicting signaling pathways involved in osteogenic cell-mediated breast cancer cell reprogramming based on MCF7 Translating Ribosome Affinity Purification (TRAP) sequencing analyzed.

**Supplementary Figure 5, related to Figure 5.**
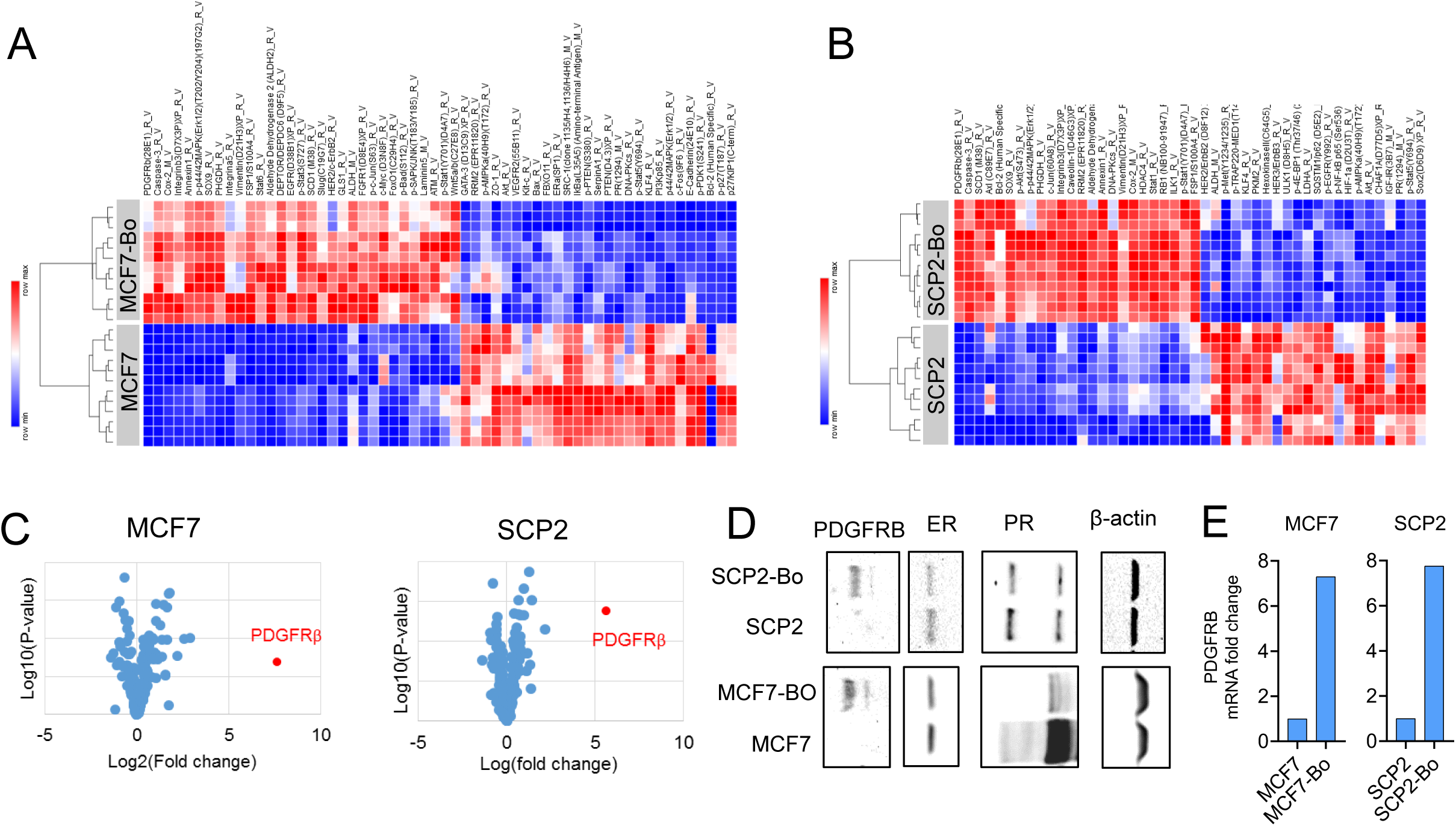
A. Heatmap showing differentially expressed proteins from RPPA data. Parental cells (MCF7) were compared to bone-entrained cells (MCF7-Bo). B. Similar to A with single cell-derived cell population 2 (SCP2) used as cells. Parental cells (SCP2) were compared to bone-entrained cells (SCP2-Bo). C. Volcano plot indicating protein distribution in MCF7-Bo and SCP2-Bo cells relatively to parental cells (MCF7 and SCP2) based on expression fold change (Log2) and p-value (Log10). D. Immunoblotting showing the expression of PDGFRB, ER, and PR in parental (MCF7 and SCP2) and bone-entrained cells (MCF7-Bo and SCP2-Bo). E. Quantitative PCR showing changes in PDGFRB expression between parental and bone-educated MCF7 and SCP2 cells. (n= 3 technical replicates)

**Supplementary Figure 6, related to Figure 6:**
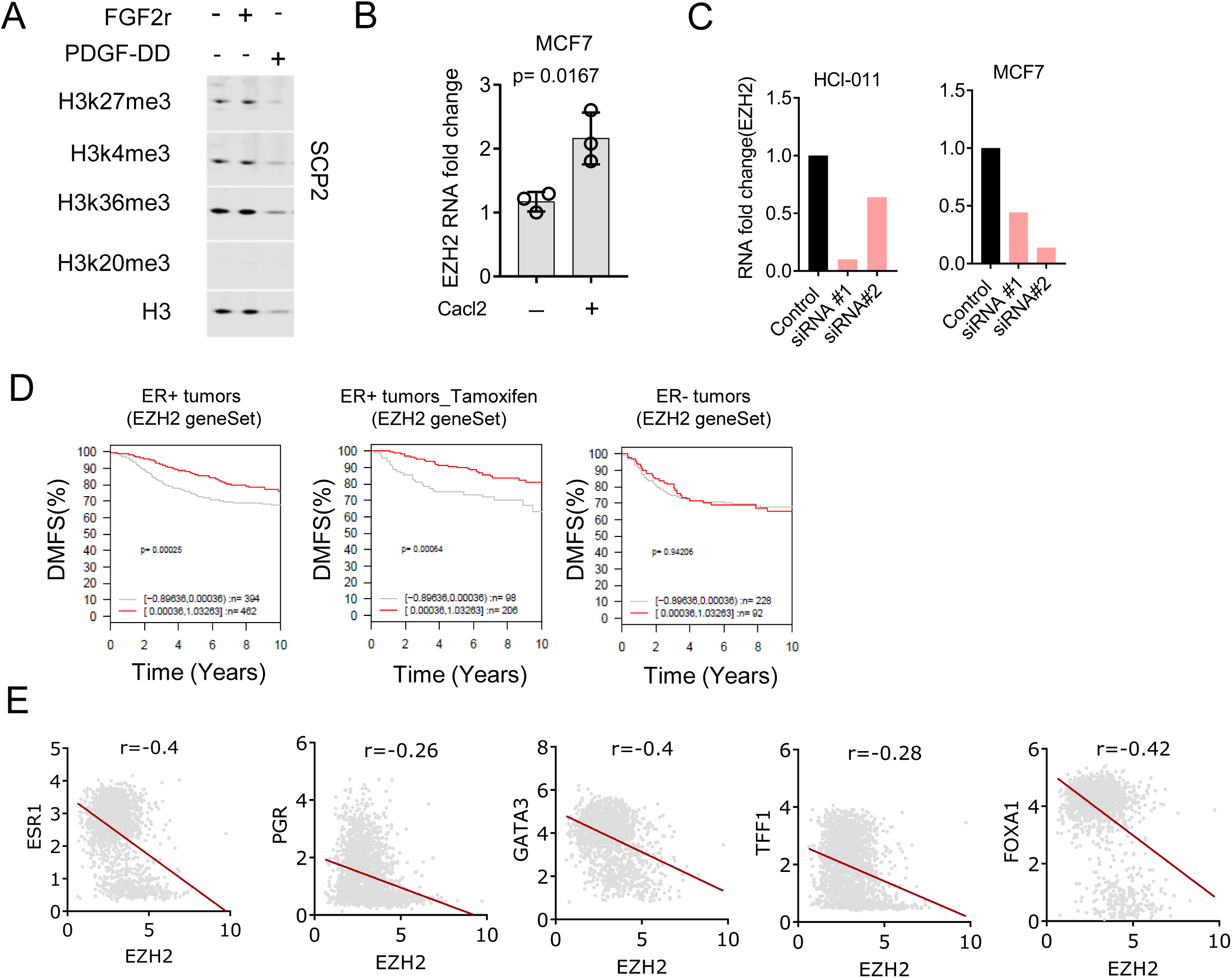
A. Immunoblotting showing the effect of FGF2 and PDGF-DD recombinant on histone modifications in SCP2. Cells were treated with 20ng/ml recombinant for 24h. B. Quantitative PCR showing the promoting effect of Calcium on EZH2 expression in MCF7 cells (n= 3 independent experiments). C. Quantitative PCR showing knockdown effect of 2 different siRNAs (s4916 and s4918) on EZH2 after 72h. MCF7 cells and HCI011 primary cells were used (n= 3 replicates). D. EZH2 target genes correlate with distant metastasis-free survival in patients with ER-positive, but not in ER-negative, breast cancer. Survival curves were generated from GOBO database (http://co.bmc.lu.se/gobo/gsa.pl) after inputting EZH2 target gene set. E. Scatter plots showing negative correlations between EZH2 and luminal markers in breast cancer patients. Gene expression was extracted from METABRIC dataset (http://www.cbioportal.org<u>./</u>). All correlations are significant (n=1866 and p-values < 10^-5^).

**Supplementary Figure 7, related to Figure 7:**
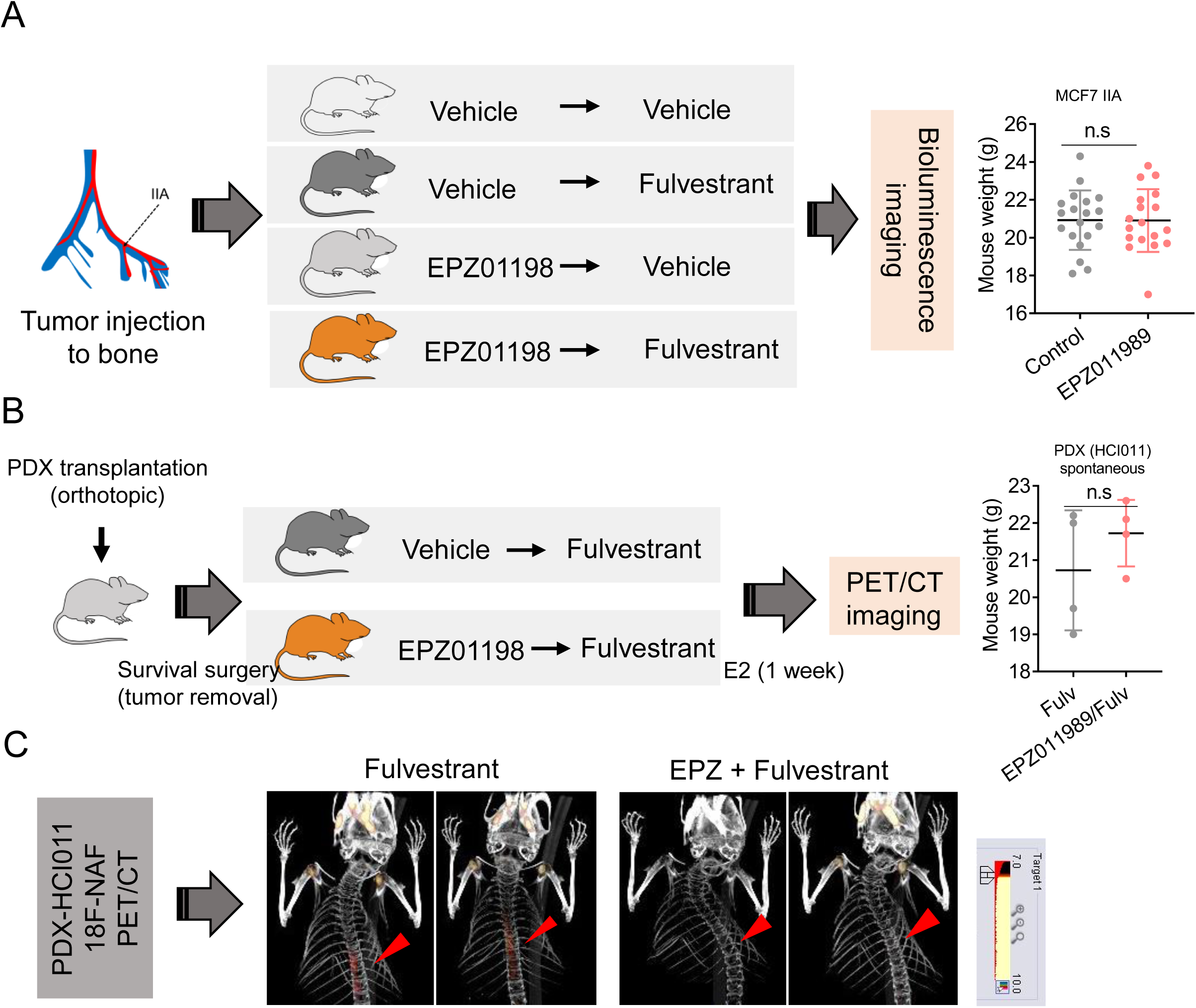
A. Diagram showing experimental procedures and treatment conditions used to assess therapeutic effects of EZH2 inhibitor (EPZ011989) in combination with fulvestrant on MCF7 bone metastasis. Tumor burden was acquired by bioluminescence (BLI). Dot plot show similar weight between vehicle and EZP011989 group after 3 weeks of treatment (n= 19 mice per group). *n.s: non-significant (P value <0.05)* B. Diagram showing experimental procedures and treatment conditions used to assess therapeutic effects of EZH2 inhibitor (EPZ011989) on spontaneous metastases of HCI011 PDXs when used in combination with fulvestrant (Fulv). Mouse weight was assessed between both treatment groups. n= 4 mice per group. *n.s: non-significant (P value <0.05)*. C. Representative 18F-NAF PET/CT scans showing spontaneous metastasis sites (hot spot) in bone following fulvestrant monotherapy or combination (mice pre-treated for 3 weeks with EPZ011989). Hot spots (red arrows) show enrichment of radiolabeled 18F-Sodium Fluoride (18F-NAF).

## STAR Methods

### CONTACT FOR REAGENT AND RESOURCE SHARING

Further information and requests for resources or reagents should be directed to the lead contact Dr. Xiang H.-F. Zhang at

#### Animal studies

All animal experiments were in compliance with Institutional Animal care and Use Committee of Baylor College of Medicine. Nude mice [Athymic Nude-Foxn1^nu^] and Scid/Beige mice [C.B-17/IcrHsd-Prkdc scid Lyst bg-J] were purchased from Envigo.

#### Patients derived xenografts (PDXs) and Primary cells

ER+ PDX models were kindly provided by Alana L. Welm (HCI011) and Matthew Ellis (WHIM9). All PDXs were maintained in SCID/Beige mice. For mechanistic studies, ER+ primary cell lines were generated from HCI011 and maintained in RPMI media supplemented with 10% FBS, 1X antibiotics Penicillin/Streptomycin and 1X antimycotics Amphotericin (Gibco#15240062).

HCI011 cells easily form clusters and tend to grow as mammospheres. To generate HCI011 primary cell lines in 2D culture, we used primary tumor-derived stromal cells as bedding for the cancer cells. Media was changed every 48 hours.

#### Cell lines

Human estrogen receptor positive (ER+) breast cancer cell lines MCF7, T47D, MDA-MB-361 and ZR75-30, pre-osteoblast cells hFOB-1.19, mesenchymal stem cells MSC, pre-osteoclast U937 and the mouse pre-osteoblast MC3T3-E1 were purchased from *American Type Culture Collection (ATCC). The Human ER+ breast cancer cell line* ZR75-1 was kindly provided by Dr. Rachel Schiff. MCF7 sub-clonal populations SCP1, SCP2, SCP3, and SCP4 were generated from single clones of parental MCF7 cells.

#### Oophorectomy

Mouse ovaries were removed using the previously described surgical procedure (Ström et al., 2012). Briefly, mice were anaesthetized with 2% isoflurane, and placed on a temperature-regulated heat pad. The dorsal area covering the lumbar vertebrae was shaved to display a 3x3 cm and disinfected. A 1 cm mid incision was performed on the skin and a 0.5 cm incision in the peritoneum allowed access to the either ovary. Ovary was removed and peritoneum closed using an absorbable suture (Ethicon Vicryl #J497G). Skin closure was completed using EZ clips (Stoelting # 59027) and mice were monitored for recovery.

#### Intra-Iliac-Artery (IIA) and mammary fat pad (MFP) injections

Intra-iliac-artery (IIA) injection was performed as previously described(Yu et al., 2016). Breast cancer cells were trypsinized, pelleted, washed twice with PBS, collected in cold PBS and kept on ice. For established breast cancer cell lines MCF7 and ZR75, 5×10^5^ cells were injected into the internal iliac artery to generate bone metastasis. For PDX models, a maximum of 200,000 cells were injected. Mammary fat pad (MFP) injections was performed as previously described (Zhang et al., 2019). Cells were prepared as for IIA injection. In cases were both IIA and MFP injections were conducted on the same host, only half of the total cell number was injected per site (50% in bone and 50% in mammary fat pad). Mice were provided with drink water containing 8µg/ml 17-beta-estradiol(Levin-Allerhand et al., 2003; Welsch et al., 1981) to increase tumor take rate.

#### Survival surgery for PET Imaging

Animals were transplanted with fresh pieces of tumor (∼1×1 mm) or 5×10^5^ cells from dissociated HCI011 PDXs tumor. Estrogen was provided through drink water or pellet implantation to increase the tumor take rate. When tumors reached 1×1 cm, a survival surgery was performed to remove primary tumors and remaining estrogen pellets. Estrogen supplementation of drink water was also stopped. Wounds were left to cure for 2 weeks followed by a 3-week vehicle or EPZ011989 treatment (oral gavage; 125mg/kg; twice daily). A day after the last EPZ011989 treatment, 250mg/kg of fulvestrant was injected to each mouse through subcutaneous injection, once a week for 2 consecutive weeks. To assess treatment efficiency, all mice were exposed to estrogen-supplemented drink water for a week before 18F-Fluorodeoxyglucose positron emission tomography (PET) and computed tomography (CT) scans were performed.

#### Drug treatments

EPZ-110989 was kindly provided by Epizyme and stock solutions were prepared following the company’s recommendations, using 0.5% Sodium carboxymethyl cellulose (NaCMC) and 0.1% Tween-80 as vehicle. A dosage of 125 mg/kg of EPZ-011989 or vehicle was administered twice daily by oral gavage (200ul) for 21 days. Fulvestrant was solubilized in 5% DMSO and 95% corn oil and administered subcutaneously at 250 mg/kg per mouse, once a week for 2 consecutive weeks.

For 3D cocultures and bone-in-culture array (BICA) experiments, 1µM of GAP19 (cat#5353) - a gap junction (CX43) inhibitor, and 1µM of FK506 known as Tacrolimus (cat#S5003) – a calcineurin inhibitor, were used to block calcium signaling. Similarly, 2.5 µM BGJ398 and 10uM Sunitinib were used to inhibit FGF receptors and PDGFRB, respectively. Co-cultures and BICA models were incubated for 24h to allow mammosphere to form for the former and cell to attach to the bone matrix for the latter, before any drug treatment was started.

### METHOD DETAILS

#### Immunoblotting assays

Proteins extraction was performed using RIPA buffer as previously described(Rajapaksa et al., 2015). Proteins electrophoresis and transfer were performed using the XCell SureLock and the iblot system (Invitrogen), respectively. Immunoblotting was performed by using antibodies against Estrogen Receptor α (D8H8), cytokeratin 19 (BA-17), Red Fluorescent Protein (Rockland-Fisher), and β-actin (8H10D10). Images were captured using the Odyssey system (Li-cor).

#### Tissue processing, immunohistochemistry (IF)

Tissues were processed with the help of the Breast Center Pathology core at Baylor College of Medicine. Immunofluorescence staining was performed using antibodies against human ERα (D8H8 and 6F11), EZH2 (D2C9), β-actin (8H10D10), Cytokeratin 8 (TROMA-I), α-Smooth Muscle Actin (D4K9N), Vimentin (D21H3).

#### Image acquisition and quantification

Most images were acquired with the Leica TCS SP5 or the Zeiss LSM 880 with Airyscan FAST Confocal Microscope. A 40X objectives was used to capture images used for nuclear IF quantifications. Each set of experiment was captured the same day to reduce technical biases. Samples stained apart and imaged at different time points were merged after normalizing the IF intensity by the mean expression of each cohort. Images were quantified using ImageJ 1.52i.

#### Tumor classification

Metastases were classified based on cell number. In average, micro-metastases were defined as lesions below 100 cells. The average maximum cell number from all model combined was 142 for macrometastases (Large lesions) and 87 for micrometastases (small lesions) which gives a fold change superior to > 1.5 between the two experiments stages of metastasis. The tumor size between different models (HCI011, WHIM9, MCF7 and SCP2) of bone metastasis was variable due to differences in tumor aggressiveness. Accordingly, we used the fold change (1.5 +/- 0.2) as a more consistent variable to segregate tumors into micrometastasis and macrometastasis.

#### Recombinant protein and calcium treatments

All protein recombinant experiments were performed in low serum media (2% serum). Protein recombinants for FGF2 (#130093838), PDGF-BB (100-14B), PDGF-CC (100-00CC), PDGF-DD (1159-SB-025) were diluted in PBS and used at a concentration of 20 or 100ng/ml. To evaluate the endocrine resistance attribute of FGF2 and PDGF recombinants, cells were starved for 48 hours in 2% charcoal stripped media before a 20nM fulvestrant treatment. All experiments involving cells growth were performed in 3D culture and bioluminescence was assessed 72h post treatment. For western blot, short term treatments were performed for 24 hours and long term ones were up to 72 hours. Western blot experiments were performed in 2D in most cases.

*Calcium treatment:* 2×10^6^ cells (MCF7 or ZR75-1) were cultured in regular medium for 24 hours. Regular medium was replaced with calcium-free minimum essential medium (S-MEM) and treated with vehicle or 2 mM Calcium chloride for 24h. Cells were harvested and lysate extracted to assess the effect of calcium on ER by immunoblotting.

#### Live imaging

For in vivo experiments, all cells were pre-labelled with Luciferase fused to GFP or RFP as previously described (Wang et al., 2015). 500,000 breast cancer cells were injected in bone or mammary fat pad, except if stated otherwise. Tumors growth was monitored using the IVIS Lumina II system. Briefly, mice were anesthetized in isoflurane chamber (2%) and 100 µl of D-Luciferin was administered through retro-orbital injection to each mouse before image acquisition.

For in vitro experiments, 10,000 cells were plated in low attachment 96 well plates to assess cell growth at 72 or 96 hours. For conditions demanding estrogen depleted media and starvation, 20,000 cells were cultured per well. Images were acquired after adding 1X concentration of D-luciferin containing media to each well.

#### Reverse phase protein arrays (RPPA)

MCF7 and SCP2 cell lines were injected to bone using intra-iliac artery injection. After 5 weeks of metastasis formation, bones were collected in aseptic conditions and dissociated to generate bone-entrained SCP-Bo and SCP2-Bo cell lines. Cells were cultured in DMEM 10% FBS supplemented with 1X antibiotics (penicillin, streptomycin) and antimycotics (Amphotericin). Bone-educated cells were FACS-sorted and maintained in 2D culture. Approximately 5×10^6^ non-entrained and bone-entrained MCF7 and SCP2 cells were harvested in freshly prepared RPPA lysis buffer containing protease and phosphatase inhibitors. Protein lysate was cleared twice via centrifugation (14,000g for 15min at 4°C). A BCA assay was adopted for protein quantification (ThermoFisher #23225). All samples were diluted in RPPA solution and SDS to a final concentration of 0.5mg/ml and heated for 8min at 100°C for protein denaturation. RPPA was performed as previously described (Welte et al., 2016). In brief, samples and control lysates were spotted onto nitrocellulose-coated slides (Grace Bio-labs; array format of 960 lysates/slide or 2880 spots/slide). The automated slide stainer Autolink 48 (Dako) was used to probe 236 antibodies (against total and phosphor-proteins) on slides. Control slides were incubated with antibody diluent. A biotinylated secondary antibody was probed by streptavidin-conjugated IRDye680 fluorophore (LI-COR Biosciences) and total protein was detected with Sypro Ruby Protein Blot Stain according to the manufacturer’s instructions (Molecular Probes). All slides were scanned on a GenePix 4400 AL scanner and images were analyzed with GenePix Pro 7.0 (Molecular Devices). Samples were normalized as previously described (Chang et al., 2015). After quality control 233 antibodies remained and were used for subsequent data analysis. (Supplementary Table 1).

#### PET/CT Imaging and Analysis

*Radiopharmaceuticals and Small-Animal PET-CT:* Flourine-18 labeled fluorodeoxyglucose (18F-FDG), fluoroestradiol (18F-FES), and sodium fluoride (18F-NaF) was purchased from (Cyclotope, Houston, TX). All CT and PET images were acquired using an Inveon scanner (Siemens AG, Knoxville, TN). The mice were injected with 9.25 MBq (250 µCi) of FES and 11.1 MBq (300 µCi) of either 18F-FDG or 18F-NaF radiotracers at any given time. To identify skeletal metastases or measure tumor metabolic activities, 18F-NaF or 18F-FDG were injected intra peritoneally, and to measure estrogen activity 18F-FES was injected intravenously via tail vein. Before 18F-FDG administration, the mice were fasted for approximately 12 hours. PET and CT was performed one hour after injection of radioisotopes. During imaging, a respiratory pad was placed under the abdomen of the animal to monitor respiration (Biovet, Newark, NJ). Mice were anesthetized with isoflurane gas (1-3%) mixed with oxygen at a flow rate of 0.5-1 L/minute, and adjusted accordingly during imaging to maintain normal breathing rates. A CT scan was acquired with the following specifications: 220 acquired projections except for the 18F-NaF imaging which was 360 full scan. Each projection was 290 ms with x-ray tube voltage and current set at 60 kVp and 500 µA, respectively. A 30 minute PET scan was immediately acquired afterward. The PET scans were reconstructed using OSEM3D reconstruction method and registered to the CT scan for attenuation correction. *PET Image Analysis:* The PET images were quantified using Inveon Research Workspace IRW (IRW, Siemens AG, Knoxville, TN). Using the reconstructed PET scan, bone (hind limbs) and mammary fat pads were manually selected to form regions of interest (ROI) on the PET-CT images. Activity measurements (Bq/cm3) were divided by the decay-corrected injected dose (Bq) and multiplied by 100 to calculate tissue uptake index represented as percentage injected dose per gram of tissue. The data was represented as standardized uptake value (SUV) normalized to body weight. For PDX spontaneous metastasis to bone, a 90% SUVmax thresholding was applied to ROI.

#### Mammosphere and coculture assays

5×10^5^ cells were plated in low attachment 6 well plates (Greiner) using regular (10% FBS) of serum free DMEM/F12 media supplemented with 3% dextran-coated Charcoal stripped. For cocultures, a 1:1 ratio was used for each cell line. Cells were collected after 48 hours of culture for RNA, or protein extraction and analysis. For immunofluorescence, cells were collected in 2% cold PFA for 24 hours, washed 3 times with PBS, embedded in paraffin, and sectioned for imaging.

#### Quantitative real-time PCR

Total RNA was extracted using the Direct-zol Zymo according to the manufacturer protocol. Copy DNA was synthesized using the iScript cDNA Synthesis Kit (Biorad). All primers are indicated in supplementary table. Real-time PCR was performed on the CFX connect system (Biorad) using PowerUP SYBR Green master mix (ThermoFisher, #A25780) for amplification.

#### RNA sequencing

Library preparation was performed Nextera XT DNA library kit as described by the manufacturer (illumina). Samples were sequenced using the Nextseq 550 system (Illumina) with the help of Genomic and RNA Profiling core (GARP) at Baylor College of Medicine.

#### Translating Ribosome Affinity Purification (TRAP) sequencing (TRAPseq)

TRAP assay was adopted from previous studies(Heiman et al., 2014). Here, we stably labeled MCF7 cells with GFP-RPL10a kindly provided by Dr. William Pu from Harvard. Cells were sorted using FACSAria II to enrich for GFP-positive cells. GFP-RPL10a expressing MCF-7 cells were maintained in 2% charcoal stripped medium for 48 hours upon which, 1 million cells were seeded in 100 mm low attachment plates (Corning, cat #05-539-101) alone or in coculture with human mesenchymal stem cells (MSCs). Mammospheres were left to form for 24h before in 10nM 17β-estradiol, or fulvestrant or 100nM Tamoxifen treatment. Cells were collected for TRAP sequencing 24h later. Library was prepared using illumina Nextera XT Kit and paired-end sequencing was performed on a Nextseq 550 System.

#### Bioinformatics analysis

*TCGA data analysis (Correlation plot):* Gene expression data was extracted from cBioPortal (www.cbioportal.org)(Cerami et al., 2012; Gao et al., 2013). The two-tailed Pearson correlations between EZH2 and luminal cancer cell marker genes (ESR1; GATA3, FOXA1, PGR, TFF1) was perform using GraphPad.

#### Quantification and Statistical Analysis

Statistical analysis of single cell IF images was performed using a two-tailed unpaired Student’s *t* test. A paired Student’s *t* test or two-way ANOVA was used for all graphs involving multiple cell lines and *in vivo* experiments unless otherwise specified. For PET imaging, Mann Whitney *U*-test was used for statistical analysis. Pearson correlation was used for correlation studies. Statistical analyses that are specific to each experiment are indicated in the corresponding figure legend.

### DATA AND SOFTWARE AVAILABILITY

Data analysis was conducted in GraphPad Prism (v8.0.1) and R (version 3.4 R). Dataset were deposited in Gene Expression Omnibus(Edgar, 2002), with the following GEO accession numbers (GSE137245; GSE137270)

## KEY RESOURCES TABLE

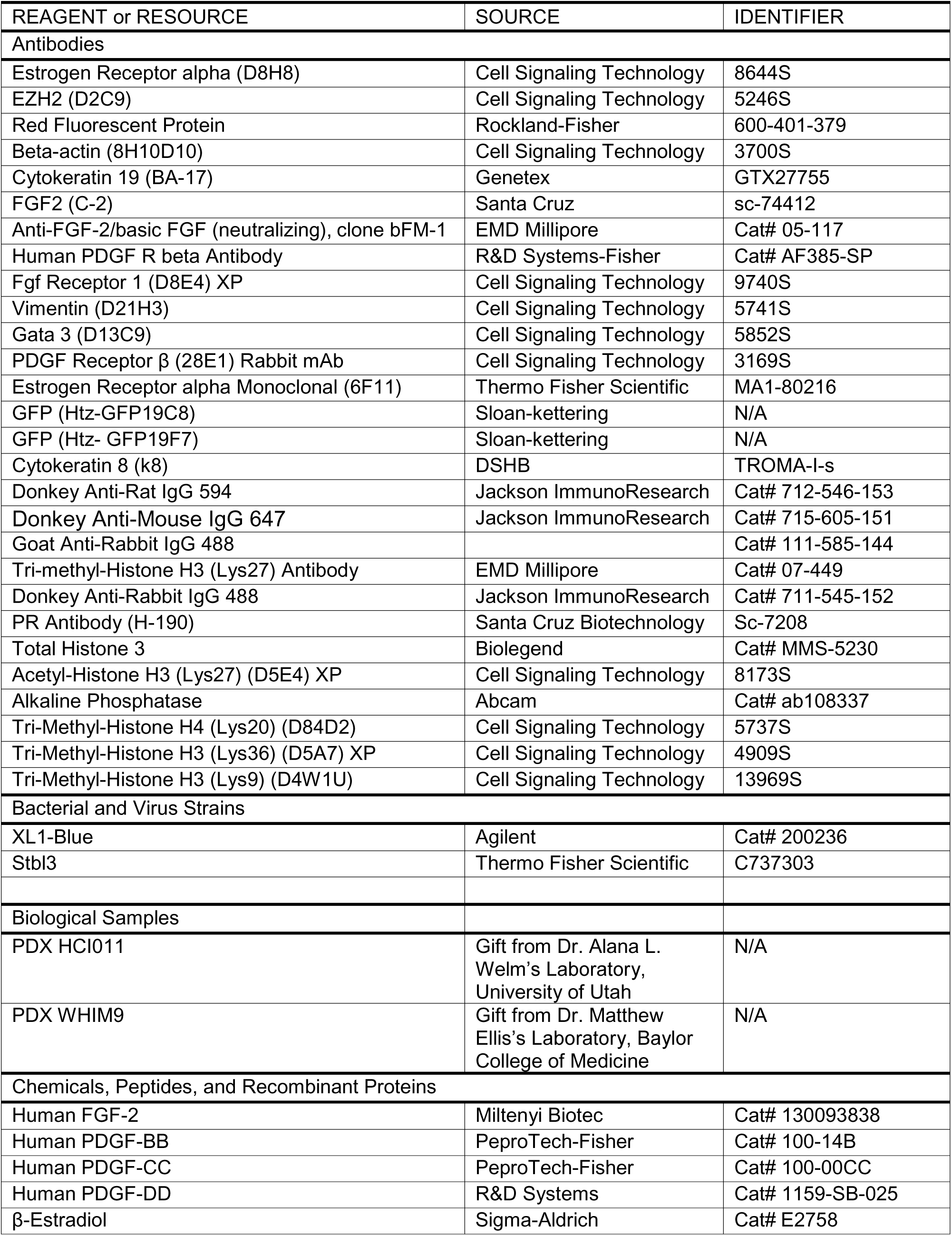
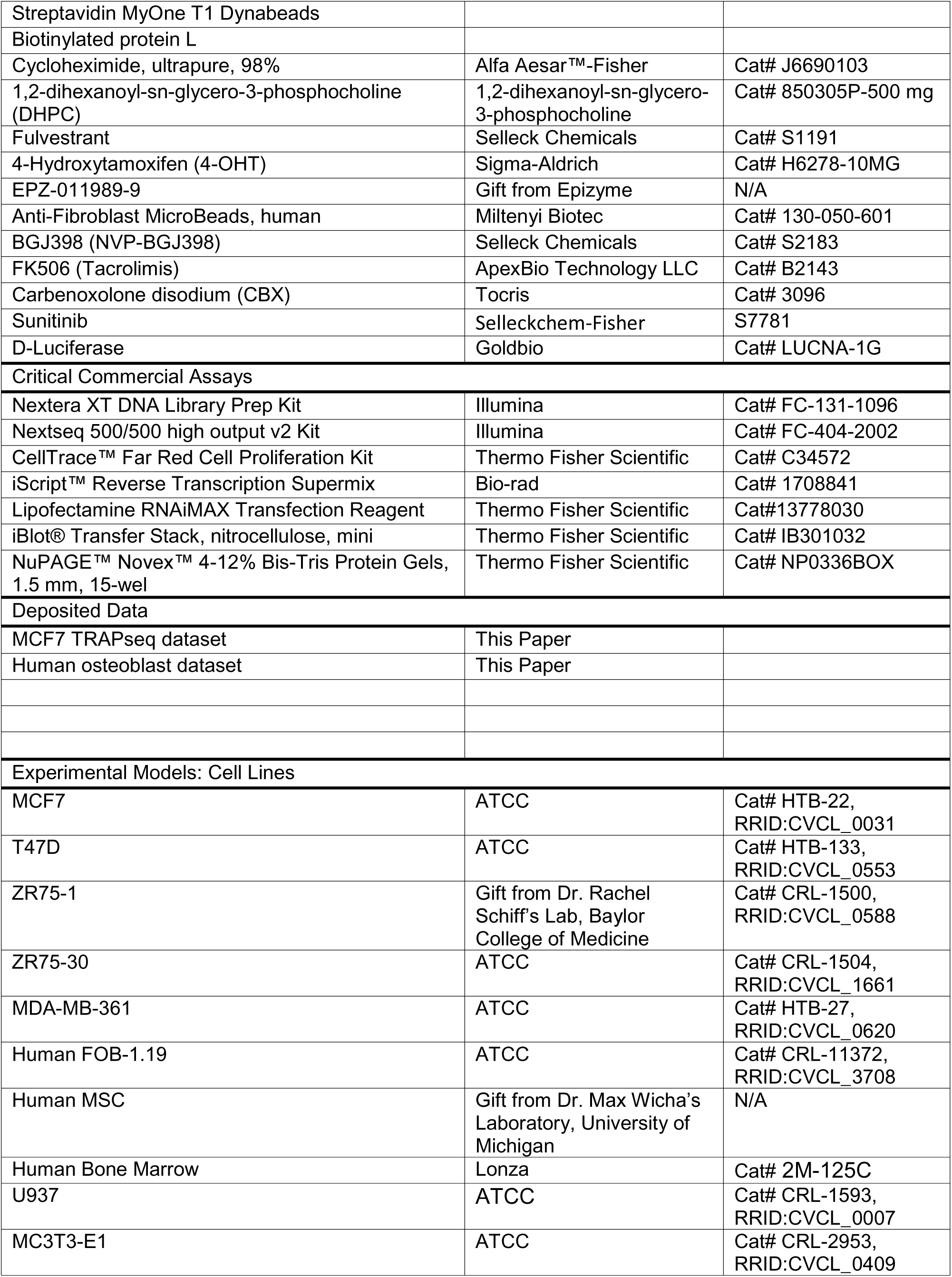
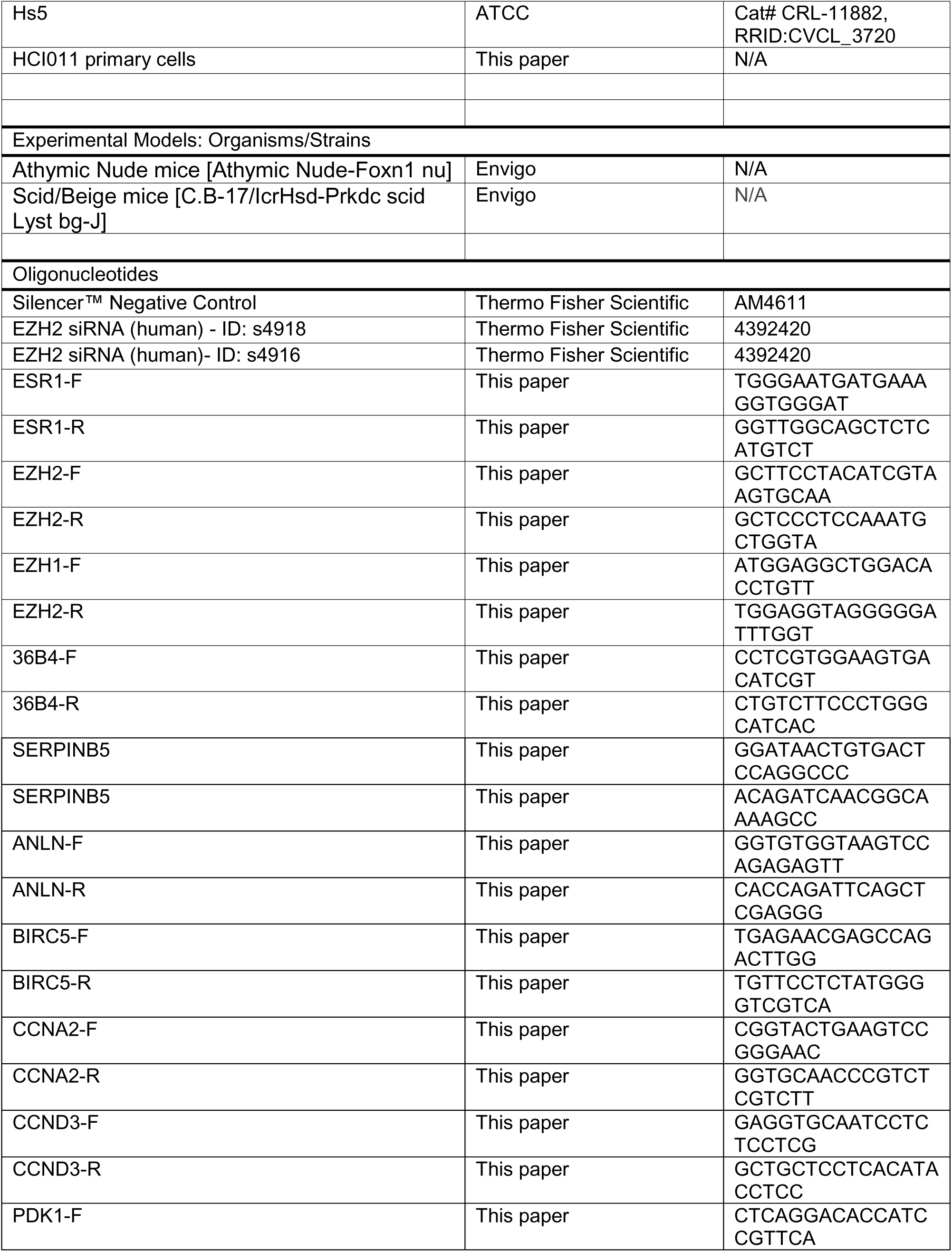
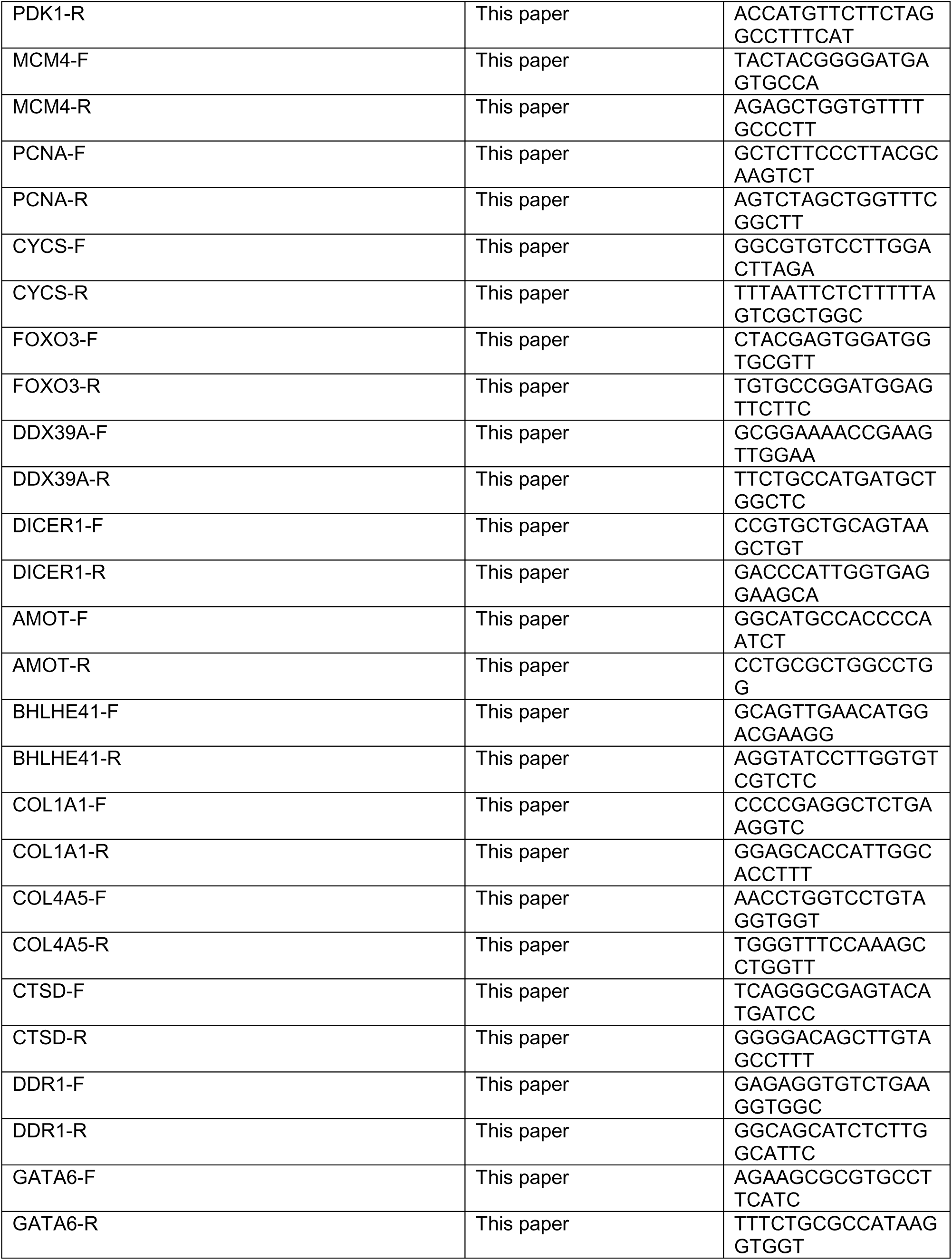
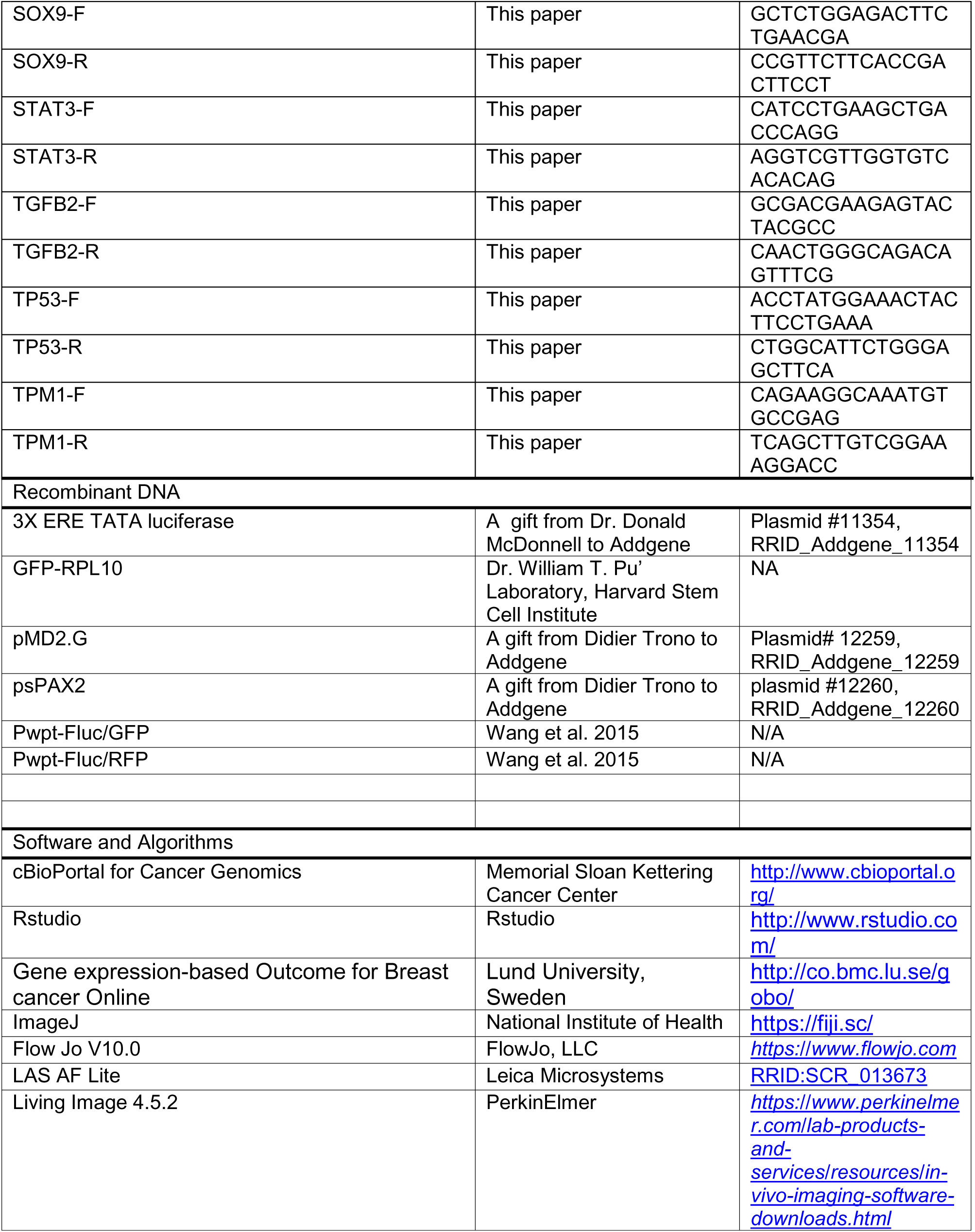
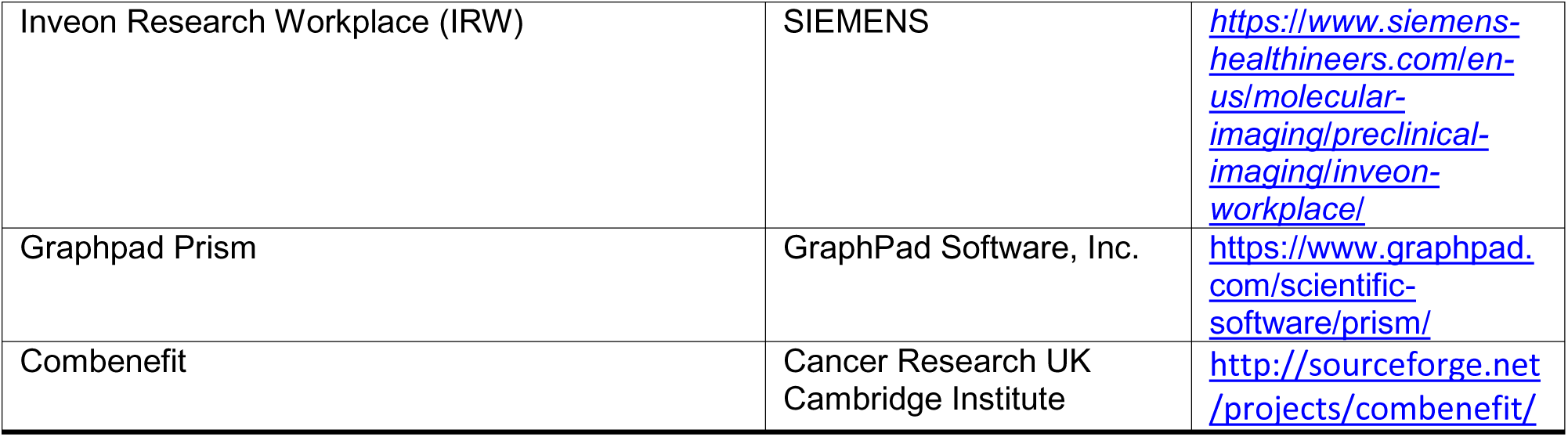

